# A synonymous mutation in *MSMEG_4729* occurs at a high frequency in spontaneous D29-resistant mutants of *Mycobacterium smegmatis*

**DOI:** 10.64898/2026.03.26.714485

**Authors:** Buhari Yusuf, Yanan Ju, Biao Zhou, Abdul Malik, Md Shah Alam, Lijie Li, Haftay Abraha Tadesse, Belachew Aweke Mulu, Cuiting Fang, Xirong Tian, Jinxing Hu, Xinyue Wang, Li Wan, Liqiang Feng, Xiaoli Xiong, Shuai Wang, Tianyu Zhang

**Author notes:** (B.Y); (Y.J.); (A.M.); (M.S.A.); (L.L.); (B.A.M.); (C.F.); (X.T.); (L.F.); (S.W.); (T.Z.); (B.Z); (X.W), (J.H).

## Abstract

Compassionate use of mycobacteriophage therapy highlights the promising potential of phage therapy as an alternative treatment option for antibiotic-resistant infections when conventional treatments fail. However, realizing the full potential of phage therapy requires addressing key challenges, including host immune responses, the limited arsenal of therapeutically-useful mycobacteriophages, and the emergence of phage resistance. Dissecting the mechanisms of phage resistance is critical for ensuring the effectiveness and sustainability of phage therapy. In this study, we demonstrate that exposure to the lytic mycobacteriophage D29 triggers diverse genetic changes in *Mycobacterium smegmatis*. A synonymous mutation in *MSMEG_4729* arises frequently but is insufficient to confer D29 resistance on its own. Instead, we identified possible Lsr2-independent activation of the lipooligosaccharide (LOS) biosynthesis cluster in a D29-resistant mutant harboring this mutation. We have also detected the possible activity of MSMEG_3213, a type II methyltransferase associated with m^6^A modifications in *M. smegmatis*. Finally, we isolated defense escape mutants (DEMs) of D29 capable of overcoming resistance in a strain with the *MSMEG_4729* synonymous mutation. This profiling of *M. smegmatis*’s likely defensive arsenal against the therapeutically-useful mycobacteriophage D29 provides a roadmap for further investigations and rational engineering of next-generation mycobacteriophages to combat drug-resistant mycobacterial infections.

**Impact statement:** Interest in phage therapy has been gaining traction recently, which is largely due to the serious threat of antimicrobial resistance. However, the efficacy and sustainability of phage therapy is threatened by certain challenges, which includes the ever existent threat of phage resistance. In this study, we identified several likely factors involved in D29 interaction with the model mycobacterium *M. smegmatis*. These findings set a roadmap for future investigations that would guide rational phage engineering to expand the currently limited arsenal of therapeutically useful mycobacteriophages as well as improve the efficiency of existing ones.

## Introduction

Antimicrobial resistance (AMR) represents a critical global health threat, driven by the rapid evolution of bacterial resistance to antibiotics. Projections suggest that AMR could cause > 39 million deaths globally between 2025 and 2050 (1). Tuberculosis, caused by *Mycobacterium tuberculosis*, remains the leading cause of death from a single infectious agent and ranks among the top 10 causes of death worldwide (2). Drug resistance further complicates tuberculosis, limiting therapeutic options and contributing to poor outcomes. Non-tuberculous mycobacteria (NTM), such as *Mycobacterium abscessus*, pose comparable or greater challenges due to intrinsic drug resistance, limited treatment regimens, and frequent therapy failure (3–7). Despite ongoing efforts to develop novel therapeutics, the persistent threat of resistance underscores the urgent need for innovative strategies to combat AMR and redefine the management of bacterial infections.

Phage therapy, the use of bacteriophages to treat bacterial infections, represents a promising alternative to antibiotic therapy. Mycobacteriophage therapy, in particular, has demonstrated safety and efficacy in managing complex mycobacterial infections, including cases refractory to conventional antibiotics. However, challenges remain in ensuring the efficient and sustainable application of phage therapy, including host immune responses and the emergence of phage resistance. Importantly, the current arsenal of therapeutically-useful mycobacteriophages is very limited, comprising only ∼10 wild-type, host-range mutants, and engineered phages with documented clinical utility against *M. abscessus*, *Mycobacterium chelonae*, *Mycobacterium avium* complex, and *Mycobacterium bovis* BCG (3–9). While host immune responses to phage therapy may vary by age (6–9), documented cases of mycobacteriophage therapy rarely report resistance evolution (3–9). Nevertheless, the threat of phage resistance exists, necessitating proactive mitigation strategies. This urgency is underscored by the identification of diverse anti-phage defense systems in mycobacteria, including in clinical isolates of *M. abscessus* (10). For example, lysogenization in *Mycobacterium smegmatis* introduces prophage-encoded defense systems that drive homotypic/heterotypic anti-phage immunity and could compromise the efficacy of phage therapy against resistant clinical isolates (10,11). Consequently, temperate phages are generally unsuitable as therapeutic agents without prior engineering to eliminate lysogenicity risks.

Additionally, disruptions in lipid homeostasis have been linked to mycobacterial phage resistance. For example, disruption of the negative transcriptional regulator-encoding gene *lsr2* by IS1096 integration results in activation of the LOS biosynthetic cluster in *M. smegmatis*, which results in membrane accumulation of phosphatidylinositol mannosides (PIMs), reduced phage adsorption and phage resistance (12). Similarly, inactivation of the *MSMEG_4242-MSMEG_4243* operon encoding the serine/threonine kinase PknL leads to elevated expression of IS1096 transposase TnpA, *lsr2* inactivation, LOS cluster activation, lipid accumulation and smooth surface morphology (13). This suggests a potential link between the *pknL*-encoding operon and the LOS-mediated phage resistance mechanism. Additionally, StpK7 (MSMEG_1200), located within the BREX-like island *MSMEG_1191-MSMEG_1200*, has been implicated in lipid homeostasis and phage resistance in *M. smegmatis* (14). PknL and StpK7 contain a single transmembrane domain each, and both function as serine/threonine-protein kinases, highlighting the potential role of this protein class in lipid metabolism and phage-bacteria interactions. Consistent with these findings, rough morphotypes of *M. abscessus* exhibit greater susceptibility to phages than their smooth counterparts, which likely stems from differences in lipid compositions of their cell envelopes (15).

The rough morphotype of *M. abscessus* is easily cleared by phages, whereas the smooth morphotype persists despite identical drug susceptibility profiles, posing a unique challenge to treatment infections involving both morphotypes (4,5,15). This dichotomy stems from fundamental differences in their lipid composition, underscoring the critical role of lipid metabolism in mycobacteria-phage interactions. In *M. smegmatis*, phage resistance has long been linked to elevated expression of *mpr,* a <u>m</u>ulti-copy <u>p</u>hage resistance gene encoding a membrane-associated exonuclease (16). Overexpressed Mpr in D29-resistant mutants is believed to cleave phage DNA thereby blocking downstream stages of D29 infection cycle (17). Notably, the DUF4352 domain of Mpr, likely of viral origin, suggests that *M. smegmatis* co-opted this protein during co-evolution with viral predators and repurposed it for defense against bacteriophages (18).

Identification of genetic factors and mechanisms underlying phage-bacteria interactions is crucial for rational phage engineering. In this study, we generated spontaneous D29-resistant *M. smegmatis* mutants and found that a synonymous mutation in MSMEG_4729, likely a putative acyltransferase encoded by the LOS biosynthetic cluster, arose frequently among the mutants. We uncovered *lsr2*-independent activation of the LOS cluster in a variant with this mutation, as well as epigenetic modifications in some mutants. We have also recently reported that exposure to D29 triggers insertion sequence (IS) rearrangements in some of these mutants, including IS6120 integration directly upstream of *mpr*, which reconstitutes a strong promoter by introducing a putative transcription factor-binding site and a canonical -35 promoter element at the integration site (19). The isolation of both D29-resistant mutants and defense escape mutants (DEMs) reveals a complex network of factors influencing D29-*M. smegmatis* interactions, hence providing actionable insights for further investigations and rational phage engineering to overcome bacterial resistance.

## Results

### Spontaneous D29 resistance in *M. smegmatis*

To generate spontaneous D29-resistant strains, *M. smegmatis* mc^2^ 155 was subjected to four rounds of sequential phage exposure. A total of 137 individual colonies were isolated and screened for D29 resistance (Figure S1), which revealed that 82.5 % (n = 113) were resistant, 15.3 % (n = 21) remained susceptible, and 2.2 % (n = 3) exhibited partial resistance (Figure 1). This high frequency of resistance underscores the ease with which *M. smegmatis* can evolve resistance under phage pressure.

**Figure 1.**
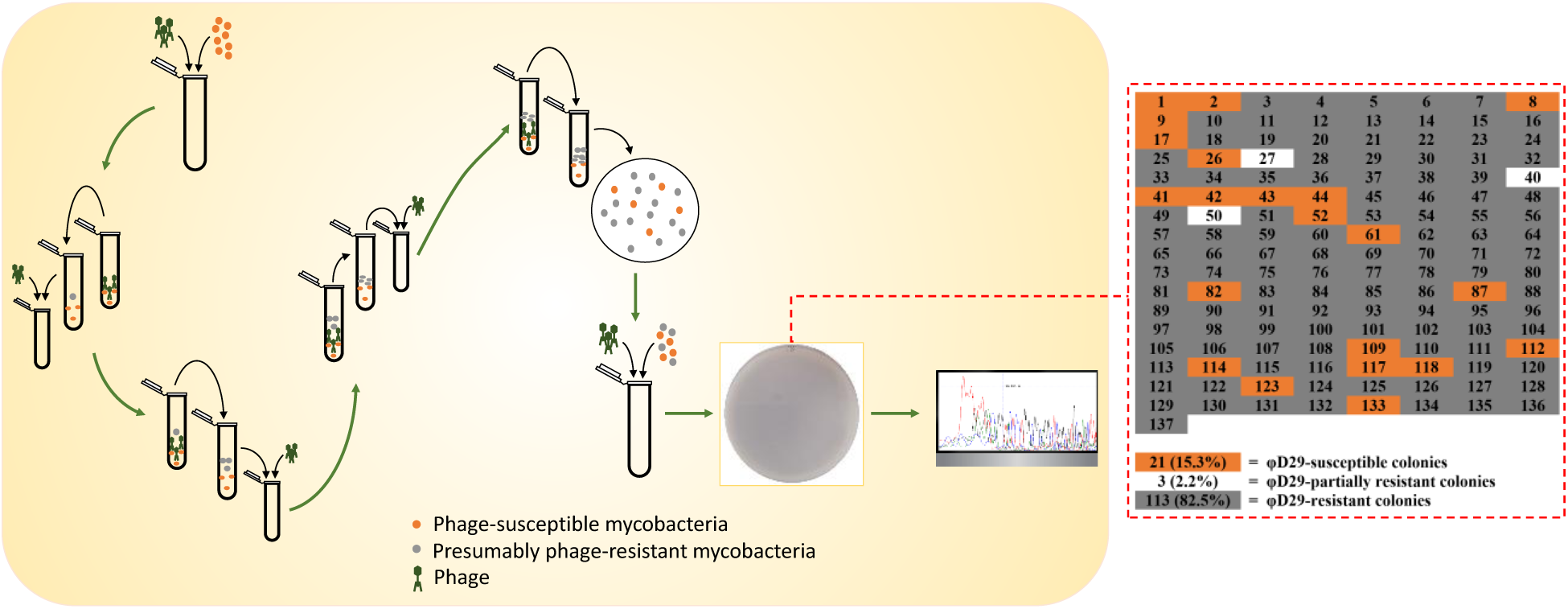
Frequency of spontaneous D29 resistance in *M. smegmatis.* To enrich for D29-resistant variants, Ms^Wt^ was subjected to four sequential rounds of extended phage exposure followed by screening of 137 single colonies. Phenotypic classification: 82.5% resistant, 2.2% partially resistant, and 15.3% susceptible.

Twenty-four D29-resistant strains were purified (Figures S2-1 to S2-3) and re-verified for resistance. All selected strains exhibited greater D29 tolerance than Ms^Wt^, with no obvious changes in colony morphology (Figure 2). While D29 forms clear zone of inhibition on Ms^Wt^, only turbid zones were observed on all resistant strains (Figure 2A), indicating reduced phage infection efficiency. Consistently, plaque assays revealed significantly fewer plaques on resistant strains compared to Ms^Wt^ (Figure 2B), with strains A72.1 and 2.6.1 showing complete resistance (no plaque formation) (Figure 2B). Notably, higher D29 concentrations partially restored susceptibility in these strains (Figure 2A), suggesting dose-dependent resistance. Phage adsorption assays indicated minimal impairment across most resistant strains, implying that resistance is not primarily mediated by inhibited adsorption (Figure 2C).

**Figure 2.**
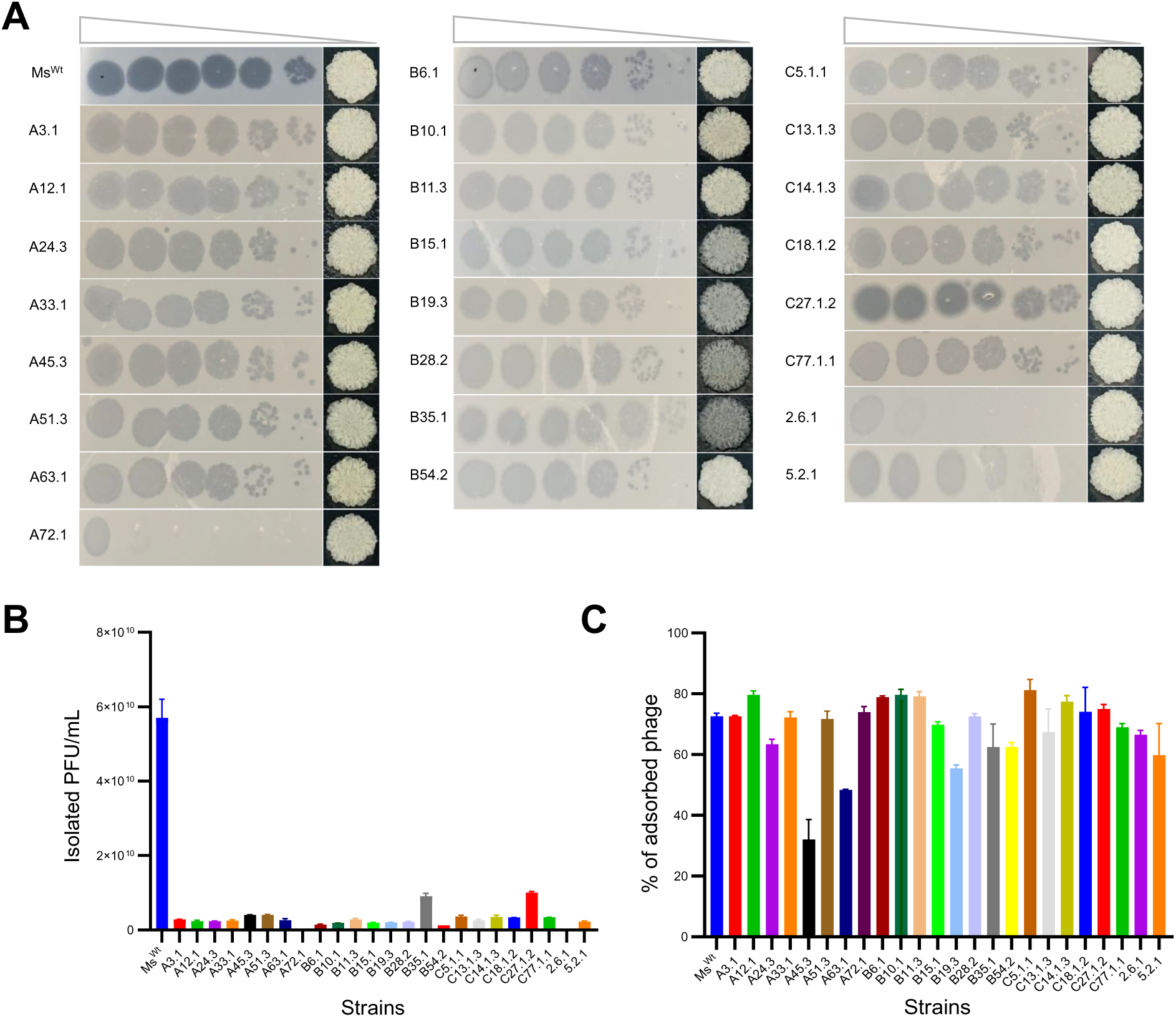
Spontaneous D29 resistance in *M. smegmatis.* Twenty-four spontaneous D29-resistant strains were purified and re-screened for D29 resistance prior to WGS. A, all selected strains were tested for D29 resistance by spot-kill assay, and all appear to show lesser level of susceptibility to D29 compared to Ms^Wt^. This is evident in the turbid zones of growth inhibition formed by D29 on the D29-resistant strains as opposed to clear zones formed on Ms^Wt^. A72.1 and 2.6.1 appear to be particularly most resistant. B, the selected strains were also tested for D29 resistance by plaque assay, which appears to be consistent with the spot-kill assay, particularly in the cases of A72.1 and 2.6.1. C, the selected strains were also screened for D29 adsorption. While terminal adsorption varies between strains, it doesn’t appear to be completely inhibited in any of the strains.

### Genetic variants in the mutant pool

Whole-genome sequencing (WGS) of the 24 D29-resistant strains revealed that 75% (n = 18) harbored genetic mutations, while 25% (n = 6) sustained no detectable mutations despite phenotypic resistance. The mutated strains displayed 1–4 alterations per isolate, with single or double mutations predominating (Figure 3). This heterogeneity suggests that both mutation-driven and potentially epigenetic or other mechanisms underlie D29 resistance in *M. smegmatis*.

**Figure 3.**
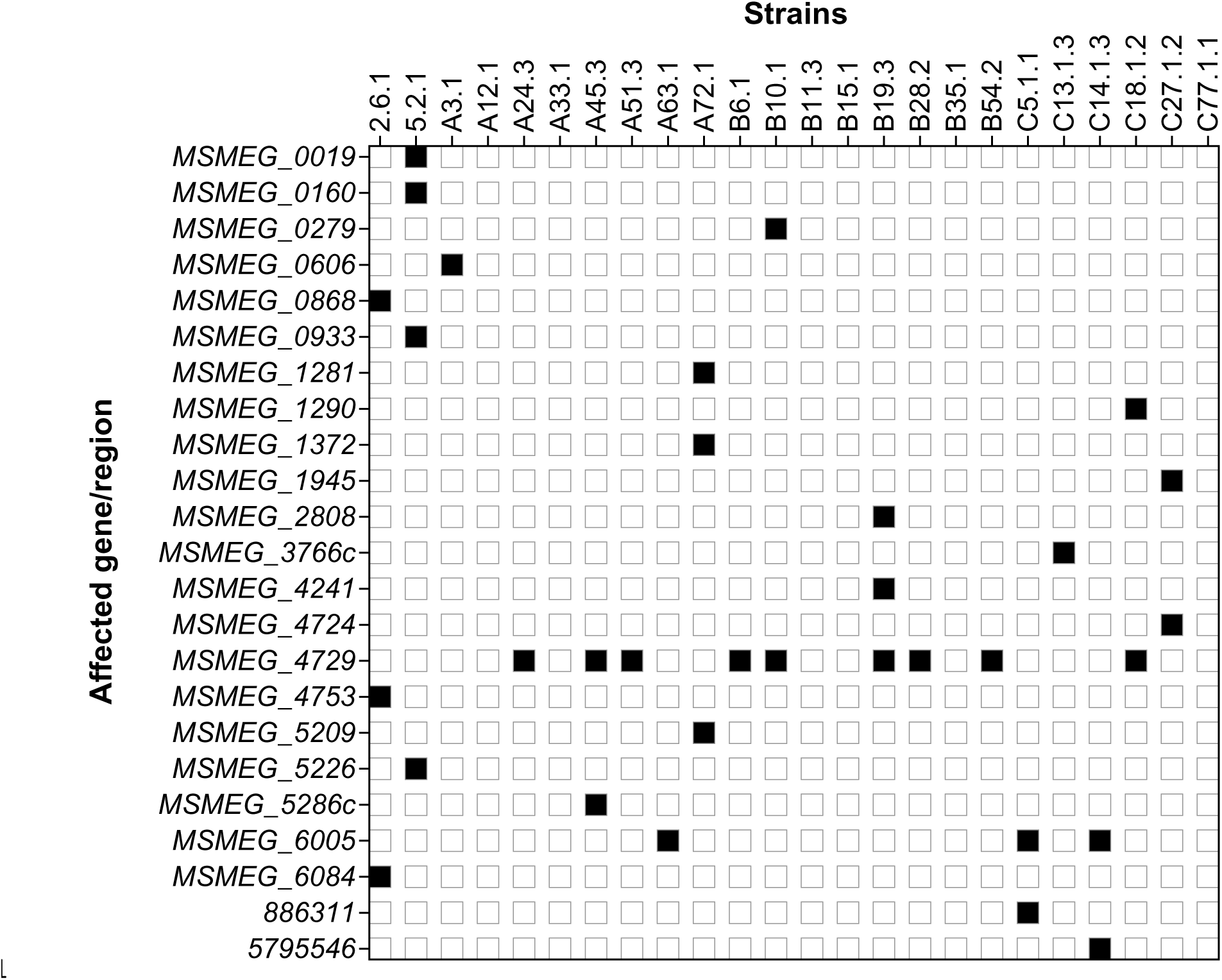
Strains and genes/regions affected by mutations. A total of 21 coding and 2 non-coding regions were affected by mutations in the D29-resistant mutants. Some mutants sustained as few as 1 and as many as four mutations. Six D29-resistant mutants are devoid of any mutation.

Mutations were detected in a total of 21 different genes and 2 non-coding regions across the D29-resistant mutants (Table 1). The affected genes spanned different metabolic pathways (Figure 4) and predominantly lie within gene clusters (Table S1). A significant proportion of these genes encode hypothetical proteins, most of which are predicted to localize to the membrane (Figure S3). Mutation types included a gene fusion, missense, frameshift, and synonymous changes (Figure S4A), with the most recurrent mutation being a synonymous alteration in *MSMEG_4729*, a hypothetical gene within the LOS biosynthesis cluster (Figure 3, Table 1, Figure S4B). This recurrence suggests a potential role for *MSMEG_4729* in phage resistance, possibly through LOS-mediated membrane remodeling.

**Figure 4.**
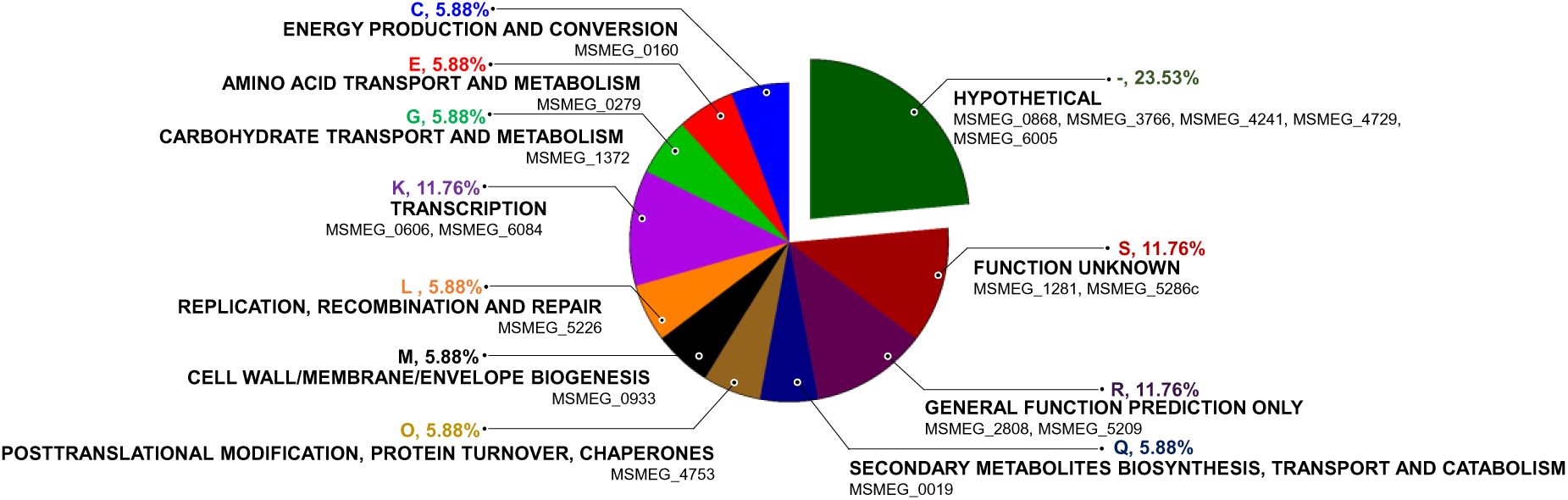
Pathway enrichment for the mutated genes. Mutated genes in the D29-resistant mutants fall into different functional categories, with most genes falling into the hypothetical proteins category, followed by those that fall into transcription, general function and unknown function categories.

**Table 1.** Genetic variants in the mutant pool.

| Position | Gene locus | Gene product | Genetic variations <sup>a</sup> |  |  |
| --- | --- | --- | --- | --- | --- |
|  |  |  | ntChange | cdChange | aaChange |
| 19157 | *MSMEG_0019* | Peptide synthetase (Amino acid adenylation) | G→A | GCG→ACG | Ala36Thr |
| 184630 | *MSMEG_0160* | Formate dehydrogenase beta subunit | A→C | GAC→GCC | Asp422Ala |
| 311327 | *MSMEG_0279* | Amino acid permease-associated region | C→G | ACC→ACG | Thr255Thr |
| 683804 | *MSMEG_0606c* | Putative transcriptional regulatory protein TetR | dupG | - | Val206fs |
| 952803 | *MSMEG_0868c* | Hypothetical protein | C→G | AAC→AAG | Asn869Lys |
| 1013795 | *MSMEG_0933* | Glycosyltransferase MshA | delC | - | Arg11fs |
| 1369672 | *MSMEG_1281* | Putative cytosolic protein | C→A | GCC→GAC | Ala513Asp |
| 1381732 | *MSMEG_1290* | Dihydroxy-acid dehydratase | C→T | CGC→CGT | Arg392Arg |
| 1472901 | *MSMEG_1372* | Sugar ABC transporter, ATP-binding protein | G→A | GGC→GAC | Gly146Asp |
| 2025189 | *MSMEG_1945* | Transmembrane cation transporter | A→C | GAC→GCC | Asp63Ala |
| 2871796 | *MSMEG_2808* | Short-chain dehydrogenase/reductase SDR | T→G | GTC→GGC | Val251Gly |
| 3831259 | *MSMEG_3766c* | Hypothetical protein | TTTCGTGCT<br>GCCAG<br>→T | - | Thr5fs |
| 4326057 | *MSMEG_4241* <sup>b</sup> | Integral membrane protein | - | - | - |
| 4817648 | *MSMEG_4724* | Oligoribonuclease | G→A | GAA→AAA | Glu176Lys |
| 4827692 | *MSMEG_4729* | Hypothetical protein | G→A | TTG→TTA | Leu219Leu |
| 4850001 | *MSMEG_4753* | Bacterioferritin comigratory protein | delG | - | Ala108fs |
| 5306182 | *MSMEG_5209* | Putative peroxidase, alpha/beta hydrolase family | dupC | - | Ser82fs |
| 5320696 | *MSMEG_5226* | Exodeoxyribonuclease 7 large subunit | TCC→ATGT | - | Ile93fs |
| 5379474 | *MSMEG_5286c* | Dihydrodipicolinate reductase | C→T | CGT→TGT | Arg112Cys |
| 6069978 | *MSMEG_6005* | Hypothetical protein | C→T | CTC→TTC | Leu31Phe |
| 6147719 | *MSMEG_6084c* <sup>c</sup> | AraC-family transcriptional regulator | A→G | GAT→GGT | Asp219Gly |
| 886311 | Non-coding | - | A→AG | - | - |
| 5795546 | Non-coding | - | G→GC | - | - |

<sup>a</sup> nt, nucleotide; cd, codon; aa, amino acid.
<sup>b</sup> *MSMEG\_4241* appears to be fused to *MSMEG\_4242* in this mutant strain by deletion of 6 bp (tgatag) from their intergenic region. There appears to be presence of three 19 bp repeats (gccacgacggcgagccgtc) that appear to be flanked and punctuated by “aatag” within this intergenic region, and these repeats do not occur anywhere else on the chromosome.
<sup>c</sup> The corresponding gene on the same chromosome (NCBI accession number CP001663) is *MSMEI\_5925c*. There appears to be another ORF (*MSMEI\_5924c*) that overlaps with this position. The nucleotide change remains the same in both *MSMEI\_5924c* and *MSMEI\_5925c*, but the aaChange in *MSMEI\_5924c* is Ile93Val (ATC→GTC). Both genes encode the same product, but *MSMEI\_5925c* is longer.

A gene fusion between *MSMEG_4241* (encoding a putative integral membrane protein) and *MSMEG_4242* (encoding a putative regulatory protein) was identified in one D29-resistant mutant. This fusion arose from a 25-bp deletion in the intergenic region, creating a mosaic sequence where two unique 19-bp repeats (gccacgacggcgagccgtc) are interspersed and flanked by three 5-bp repeats (aatag) (Figure 5). Notably, these 19-bp repeats are absent elsewhere in the *M. smegmatis* genome. Both genes reside within the same operon, indicating possible localized genetic rearrangements in phage resistance.

**Figure 5.**
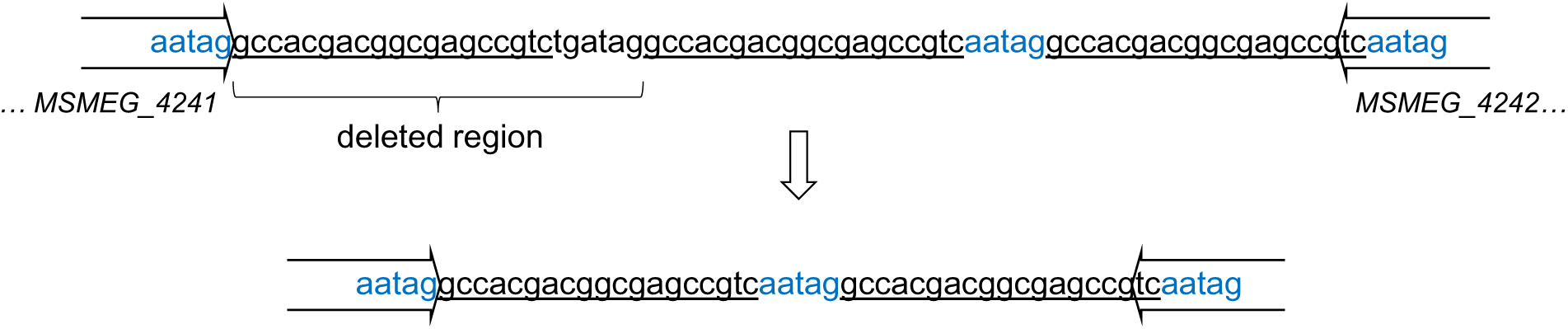
*MSMEG_4241-MSMEG_4242* fusion in the D29-resistant mutant B19.3. The D29-resistant mutant B19.3 sustained 3 mutations, including a 25-bp deletion from the non-coding region between *MSMEG_4241* and *MSMEG_4242*, resulting in what was classified as gene fusion by the variant calling program. In addition, this non-coding region contains three 19-bp sequences (underlined) that do not appear anywhere else on the entire *M. smegmatis* genome. Deletion of the 25-bp region (including one of the 19-bp sequences) resulted in a region with a mosaic appearance in B19.3, in which the remaining two 19-bp sequences appear to be separated and flanked on both sides by identical 5-nucleotide sequences (blue font).

### *MSMEG_4729G657A* mutation occurs at a high frequency in D29-resistant mutants

The synonymous mutation *G657A* in *MSMEG_4729* (*4729G657A*) was the most prevalent genetic alteration among the D29-resistant mutants (Figure S4B). To validate its prevalence, we screened all the 116 D29-resistant/partially resistant strains by Sanger sequencing and confirmed that this mutation occurs at an exceptionally high frequency (Figure 6). This high prevalence likely positions *MSMEG_4729* as a central player in D29 resistance, potentially through LOS biosynthesis-mediated membrane remodeling.

**Figure 6.**
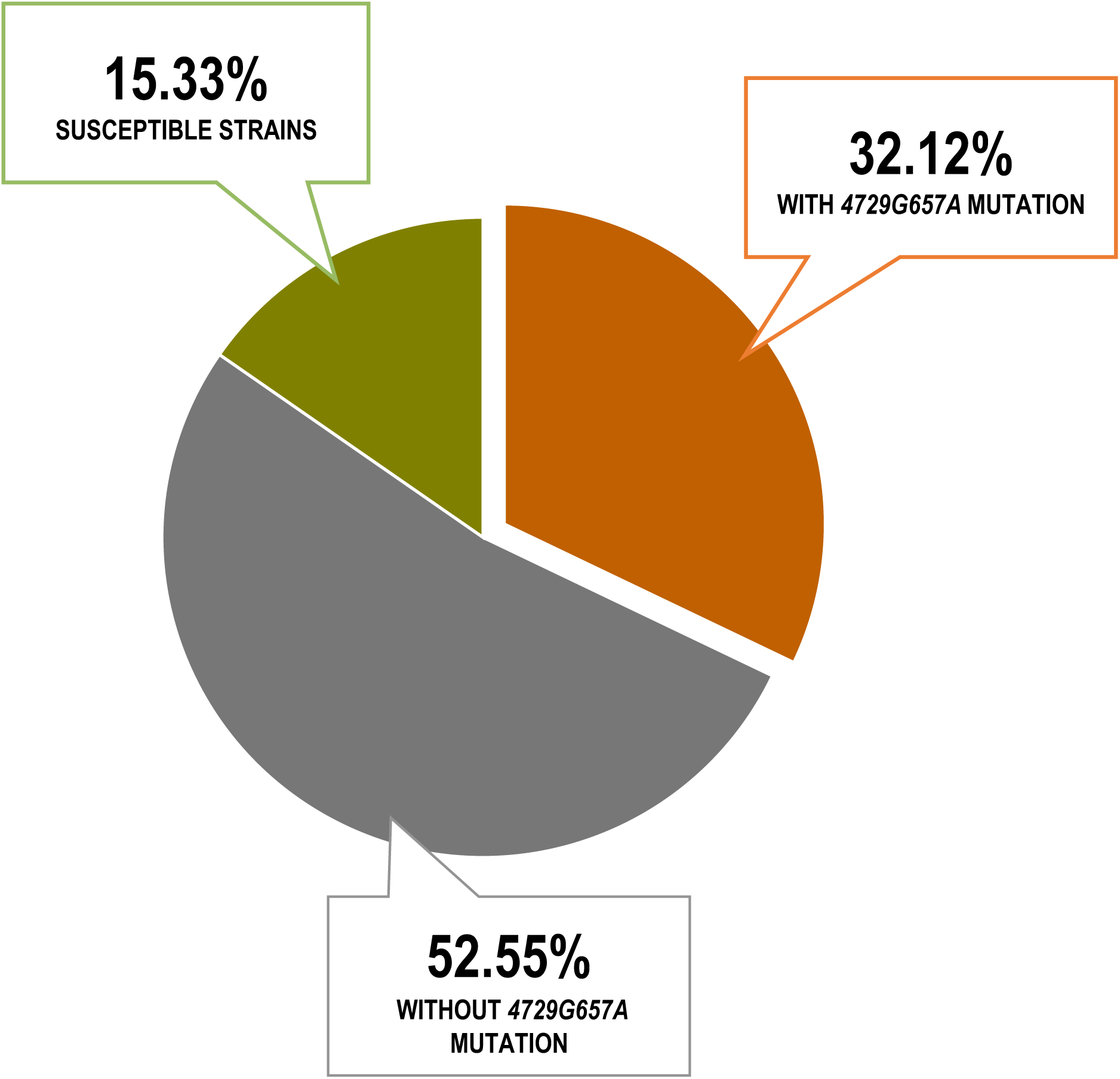
Prevalence of the *MSMEG_4729 G657A* mutation in D29-resistant isolates. A total of 137 *M. smegmatis* colonies were screened for D29 resistance (Figure S1), of which 21 (15.33 %) remained susceptible. The nucleotide substitution *MSMEG_4729G657A* led to the synonymous mutation Leu219Leu (Table 1). This mutation occurs at a high frequency across the 116 D29-resistant/partially resistant strains.

To investigate the role of *4729G657A* mutation in D29 resistance, we selected B6.1 for downstream investigations as it harbored only this mutation. Resistance was validated via spot-kill assay, plaque assay as well as growth on D29-seeded 7H10 plates. Despite bearing a synonymous substitution, often considered “innocuous”, B6.1 mutant strain exhibited robust resistance: it grew on phage-seeded plates, formed turbid (rather than clear) zones of inhibition, and produced far fewer plaques compared to Ms^Wt^ (Figure 7). To test whether D29 could adapt to this resistance, three plaques isolated from B6.1 (D29^B6.1^-I, II, III) were amplified and used to re-infect B6.1. All three variants infected B6.1 similarly to the original D29 (Figure S5), suggesting no rapid adaptation occurred during single-round passaging.

**Figure 7.**
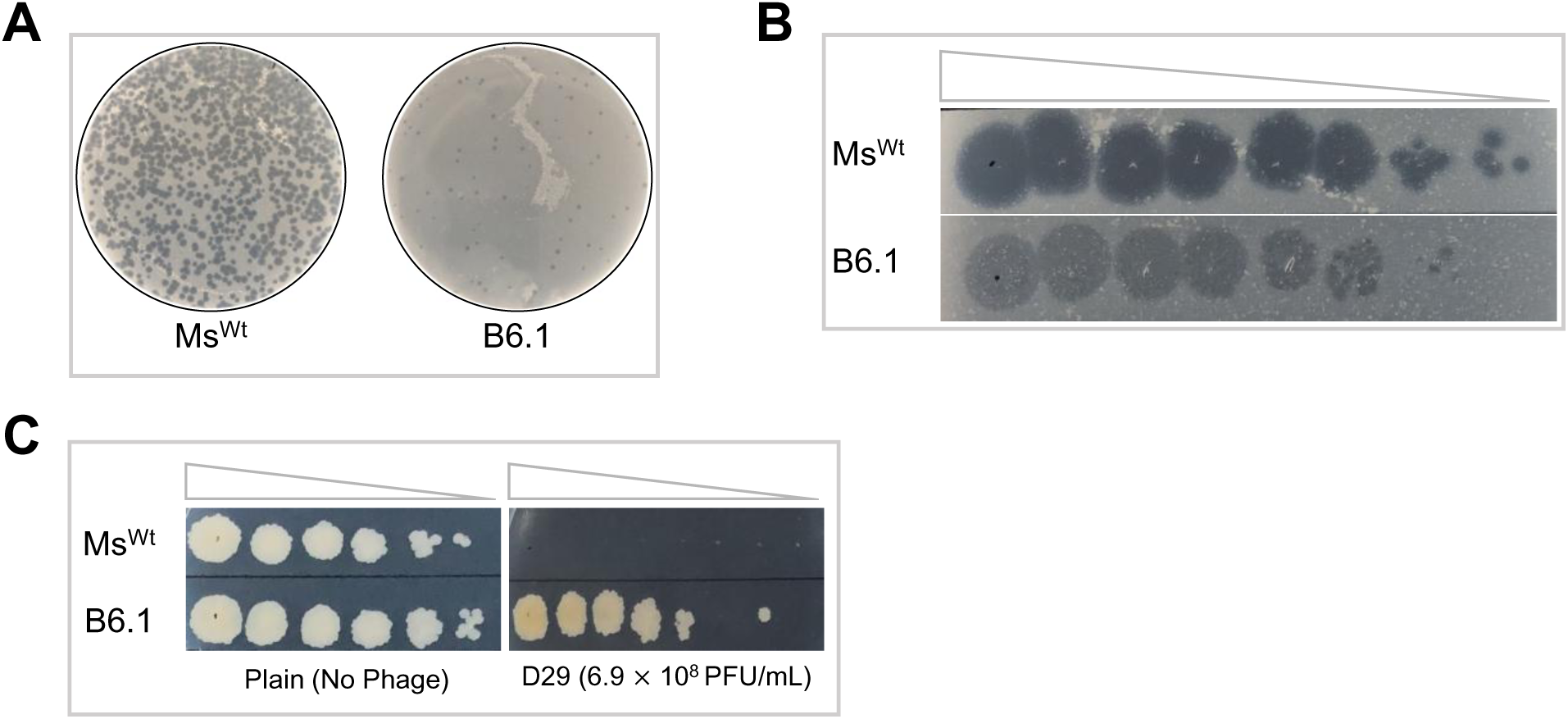
D29 susceptibility of the B6.1 mutant. B6.1 is a D29-resistant mutant that sustained only a single mutation–the *4729G657A* synonymous mutation. The strain appears to show high level of resistance to D29. A, plaque assay reveals that D29 forms fewer and rather turbid plaques on B6.1 in comparison to the numerous, clear plaques formed on Ms^Wt^. B, spot-kill assay is consistent with the plaque assay findings, confirming that D29 forms turbid zones of growth inhibition on B6.1 in contrast to the clear zones formed on Ms^Wt^. C, while Ms^Wt^ fails to grow on 7H10 plates seeded with 10^8^ PFU/mL of D29, B6.1 appears to grow well.

### *MSMEG_4729* lies in the LOS biosynthesis cluster of *M. smegmatis*

*MSMEG_4729* encodes one of the several hypothetical proteins enriched among mutations in the D29-resistant strains (Figure 4). It resides within a 15-gene cluster dedicated to LOS biosynthesis in *M. smegmatis*. Predictions from PANDA (20) indicate that MSMEG_4729 may function as an acyltransferase (Table 2) and likely localizes to the membrane (Figure S3), which may implicate it in cell envelope remodeling—a plausible mechanism for phage resistance.

**Table 2.** MSMEG_4729 function prediction using PANDA.

| Protein | Ontology | GO ID | Score | Definition (synonyms) |
| --- | --- | --- | --- | --- |
| MSMEG_4729 | BPO | GO:0046677 | 0.12 | Response to antibiotic |
|  | CCO | GO:0005886 | 0.13 | Plasma membrane |
|  |  | GO:0005737 | 0.10 | Cytoplasm |
|  | MFO | GO:0004144 | 0.06 | 1. Diacylglycerol O-acyltransferase activity (1,2-diacylglycerol acyltransferase activity<br>2. Acyl-CoA:1,2-diacylglycerol O-acyltransferase activity, diacylglycerol acyltransferase activity<br>3. Diglyceride O-acyltransferase activity<br>4. Diglyceride acyltransferase activity<br>5. Palmitoyl-CoA-sn-1,2-diacylglycerol acyltransferase activity) – catalysis of the reaction: acyl-CoA + 1,2-diacylglycerol = CoA + triacylglycerol. |
|  |  | GO:0047196 | 0.04 | 1. Long-chain-alcohol O-fatty-acyltransferase activity (wax ester synthase activity<br>2. Acyl-CoA:long-chain-alcohol O-acyltransferase activity<br>3. Wax synthase activity, wax-ester synthase activity) – catalysis of the reaction: a long-chain-alcohol + acyl-CoA = a long-chain ester + CoA. |
|  |  | GO:0016747 | 0.04 | Acyltransferase activity – catalysis of the transfer of acyl groups other than amino-acyl groups from one compound (donor) to another (acceptor). |

GO, gene ontology; BPO, biological process ontology; CCO, cellular component ontology; MFO, molecular function ontology.

The mc^2^ 155 strain of *M. smegmatis* does not produce LOS, but its parent strain ATCC 607 does (21,22). Comparative genomic analysis revealed near-identical LOS clusters in both strains (99.99% nucleotide identity), with divergence limited to MSMEG_4734 (617 aa) and its ATCC 607 ortholog, NCTC7017_04217 (555 aa) (Figure 8A). SMART domain analysis identified a transmembrane domain at positions 7–29 in MSMEG_4734—absent in NCTC7017_04217—likely due to two “C” nucleotide insertions at positions 122 and 166 (corresponding to Ser41 and Arg56) in mc² 155. The ATCC 607 LOS cluster is shorter by 2 bp (25,155 bp vs. 25,157 bp), and origin of the “C” insertions—whether biological or technical (e.g., sequencing error)—remains ambiguous (Figure 8B).

**Figure 8.**
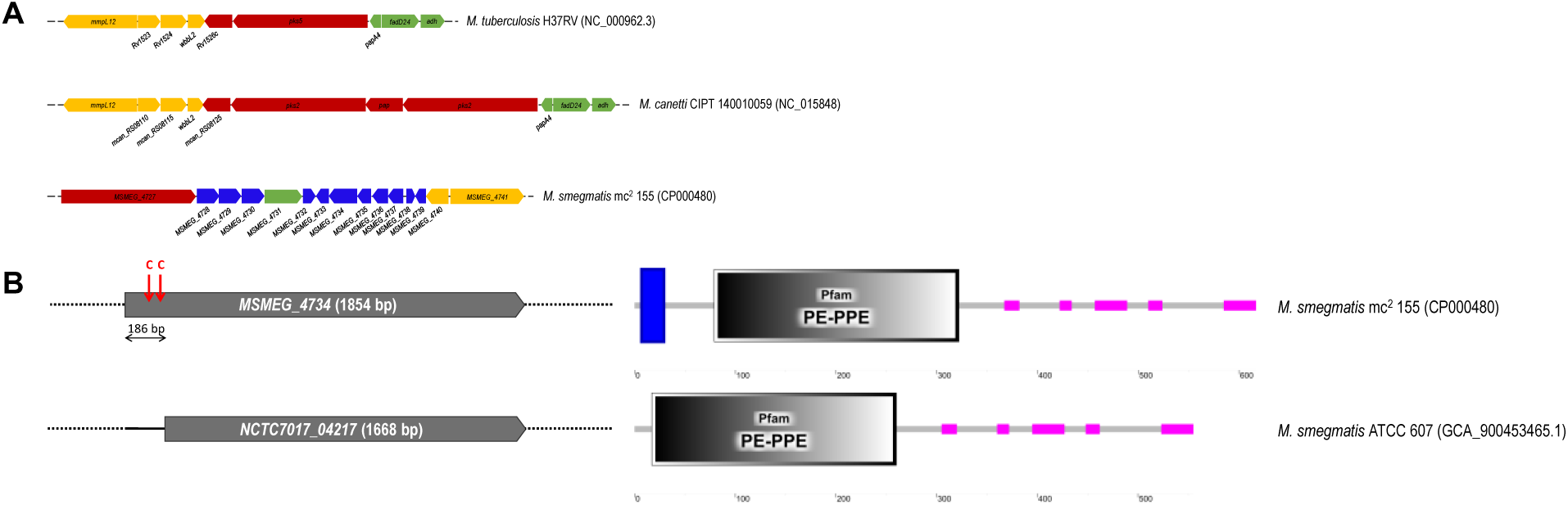
LOS cluster comparision between different mycobacterial species. Only certain species of mycobacteria produce LOS, including *M. canetti*. *M. smegmatis* ATCC 607 produces LOS, but its daughter strain–*M. smegmatis* mc^2^ 155–does not. A, cluster comparison reveals that *M. canetti* has two polyketide synthases (*pks*) in its LOS cluster–a feature known to be associated with LOS-producing *Mycobacterium* species–whereas *M. tuberculosis* and both *M. smegmatis* strains have one each. *M. tuberculosis* and *M. canetti* clusters share more locally collinear blocks (LCBs), differing only in the presence of a *pap* gene and a second *pks* gene in *M. canetti* LOS cluster. Only about 26.7 % of the *M. smegmatis* LOS genes bear similarity with those in the *M. tuberculosis* and *M. canetti* LOS clusters. Figure was drawn to scale from Mauve-generated LCBs (60). B, The LOS clusters in each of the ATCC 607 and mc^2^ 155 strains of *M. smegmatis* encode a single *pks* gene. In fact, the clusters share a 99.99 % similarity at nucleotide level. The only difference between these clusters appears to be the two “C” insertions (upper left) and a transmembrane domain (upper right) at the N-terminal of *MSMEG_4734* in mc^2^ 155. The same gene in ATCC 607 appears lack this insertion, hence making it shorter (lower left) and devoid of the transmembrane domain (lower right).

### *4729G657A* mutation may not be the sole factor responsible for D29 resistance

To establish genetic evidence for the role of *MSMEG_4729* or *4729G657A* mutation in D29 resistance, we generated a knockout strain for *MSMEG_4729* (Ms^Δ4729^) as well as an edited strain (Ms^E4729^) in which the synonymous mutation was introduced by CRISPR-Cpf1-assisted nucleotide editing (Figure S6). Ms^E4729^ was purified, and four colonies Ms^E4729-5^, Ms^E4729-19^, Ms^E4729-25^ and Ms^E4729-30^ were verified to have the desired mutation. Both Ms^Wt^ and the edited strains displayed a similar phage susceptibility profile except on D29-seeded plates (Figure S7), and adsorption doesn’t seem to be inhibited in the strains. Because the strains contain recombineering vectors pJV53-Cpf1 (with kanamycin selectable marker) and pCR-Zeo (with zeocin selectable marker), we rid the bacteria of both vectors and repeated the phage susceptibility testing. It appears that all strains lost their basal resistance phenotype after plasmid shedding (unmarking), suggesting that *4729G657A* mutation may not be the sole driving factor behind D29 resistance in B6.1 (Figures 9-10).

**Figure 9.**
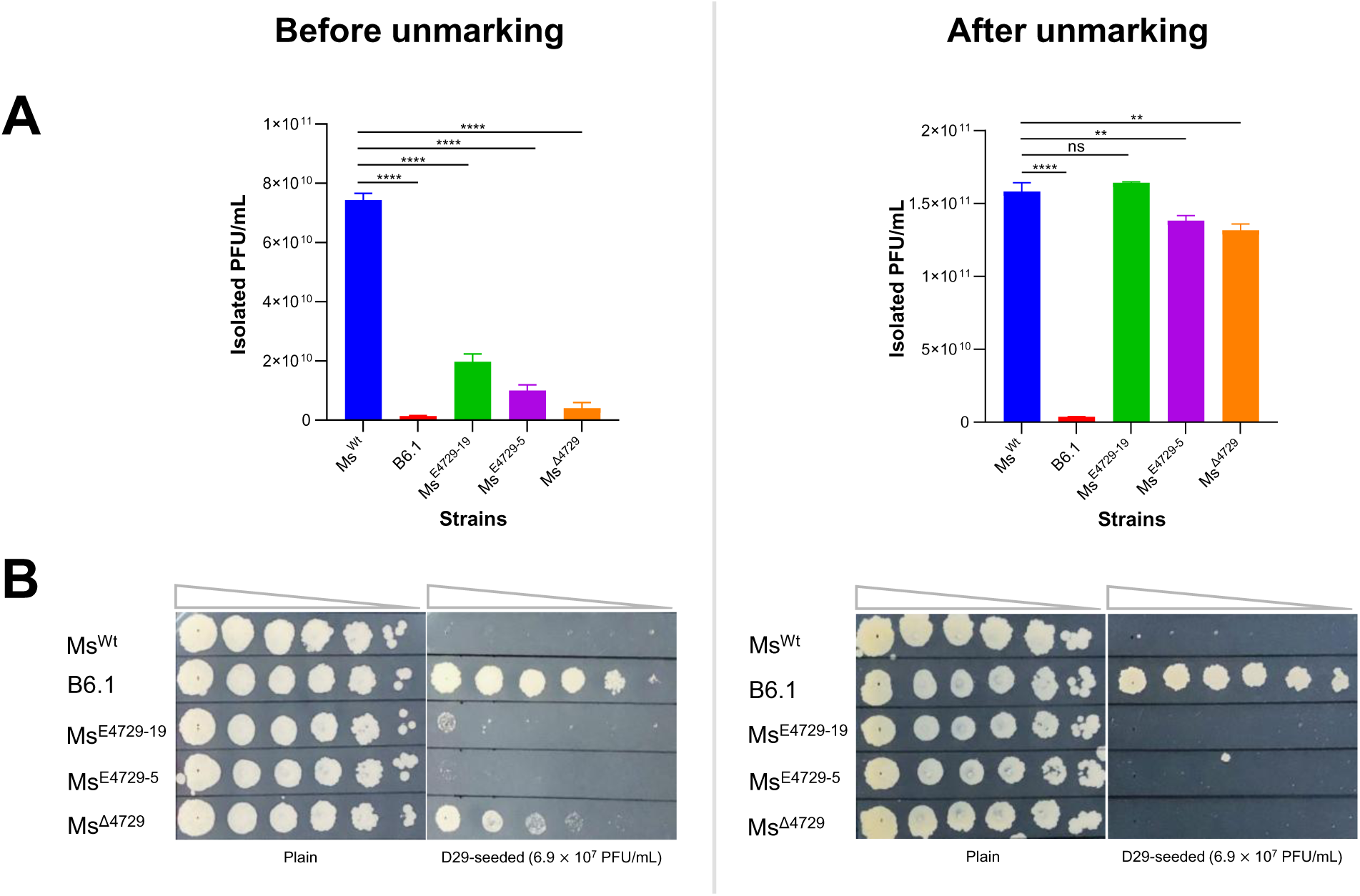
Phenotypic characterization of D29 susceptibility in engineered *M. smegmatis* strains. A, plaque assay reveals that both knockout and base-edited strains seem to show some level of resistance to D29 in contrast to Ms^Wt^, albeit to a far lesser extent than B6.1. However, both edited strains harbor recombineering vectors, and curing the strains of these vectors (unmarking) almost completely reversed their resistance to D29. B, a similar trend was seen on D29-seeded 7H10 plates, where the knockout and base-edited strains lost their basal D29 resistance after unmarking.

**Figure 10.**
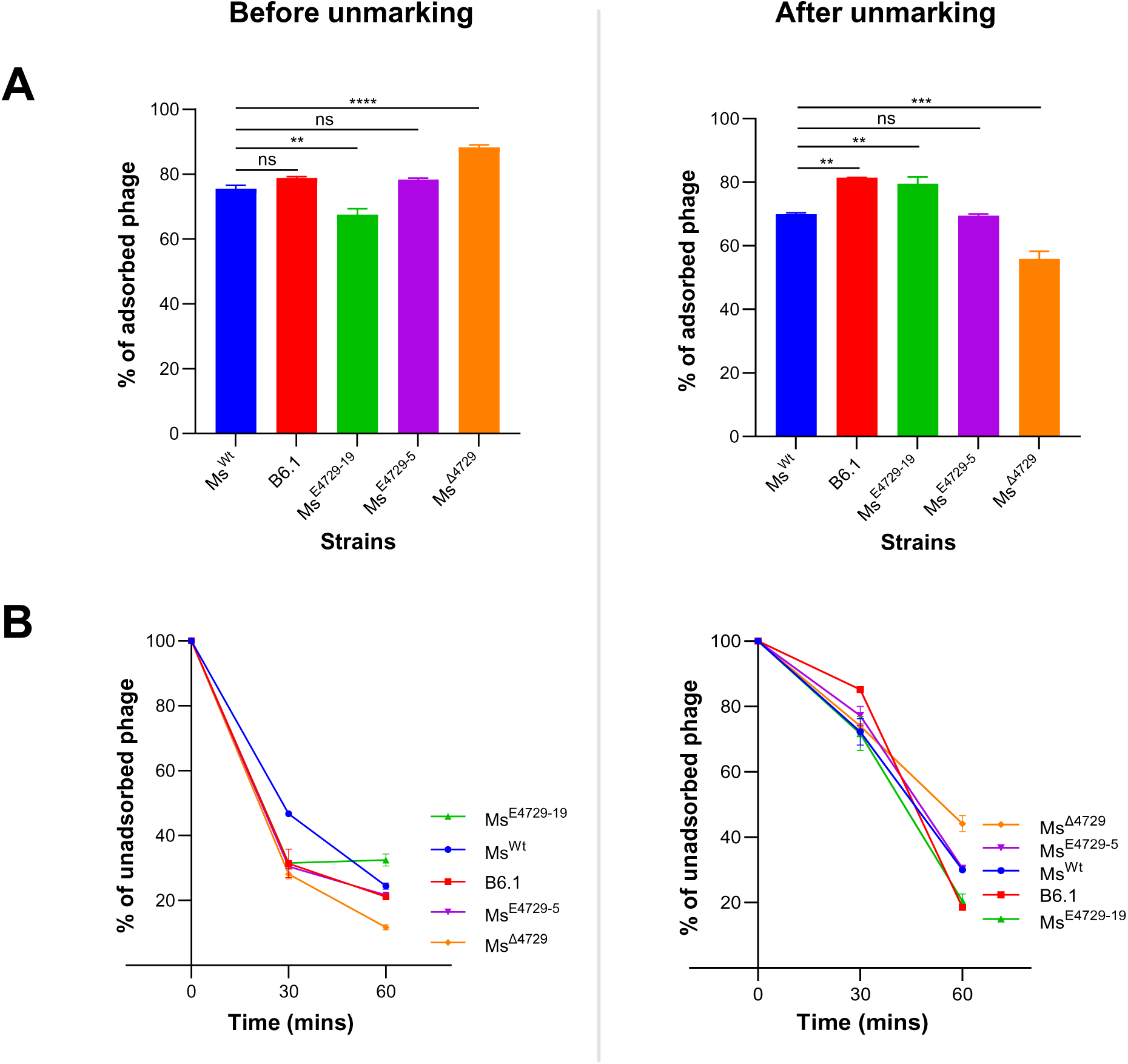
Adsorption of D29 to engineered *M. smegmatis* strains. A, terminal D29 adsorption to both knockout and base-edited strains remains similar to those of Ms^Wt^ and B6.1 and remains the same after unmarking. B, the rate of D29 adsorption to all tested strains also remained similar, but not the same. Overall, phage adsorption remains largely uninhibited in all tested strains.

We further verified this by checking growth of the following strains on D29-seeded 7H10 plates: Ms^Wt^; Ms^Wt^ harboring pJV53-Cpf1 (Ms:pJV53-Cpf1); B6.1; original unpurified edited strain (Ms^E4729^); two purified edited strains with the desired mutation (Ms^E4729-5^ and Ms^E4729-19^); and two purified edited strains without the desired mutation (Ms^E4729-1^ and Ms^E4729-2^). At the same time, all the strains were spotted on 7H10 plates containing kanamycin, zeocin or no drug (plain) to ascertain whether or not they contain either or both of the vectors. It was surprising that all strains show a greater level of tolerance than Ms^Wt^ (Figure S8), which further confirms that *4729G657A* is not the sole factor responsible for D29 resistance.

### Overexpression of *MSMEG_4729* confers *M. smegmatis* with resistance to D29

Since knockout and editing of *MSMEG_4729* did not confer D29 resistance, we overexpressed either of the wild type (*MSMEG_4729*) or mutated version (*MSMEG_4729G657A*) of the gene in Ms^Wt^ to assess their impact on phage susceptibility. Susceptibility was evaluated via spot-kill assay, plaque assay, and growth dynamics in co-culture with D29 versus uninfected controls (Figure 11). Neither overexpression strain allowed for plaque formation or clear zone of inhibition upon D29 exposure (Figure 11A, B), despite normal growth in the absence of phage. Growth in the absence of D29 infection on agar plates or in liquid medium suggest minimal differences between all strains (Figure 11C, left and middle panels). However, B6.1 and both MSMEG_4729-overexpressing strains tolerated D29 exposure right from early stages of growth, albeit to a lesser extent than B6.1 (Figure 11C, right panel), which implies that *MSMEG_4729* overexpression—regardless of the *G657A* mutation—modulates D29 resistance, likely through membrane-associated mechanisms. Therefore, while growth curves appear to show minimal differences in growth between all strains in the absence of D29, there is complete inhibition of Ms^Wt^ growth in the presence of D29.

**Figure 11.**
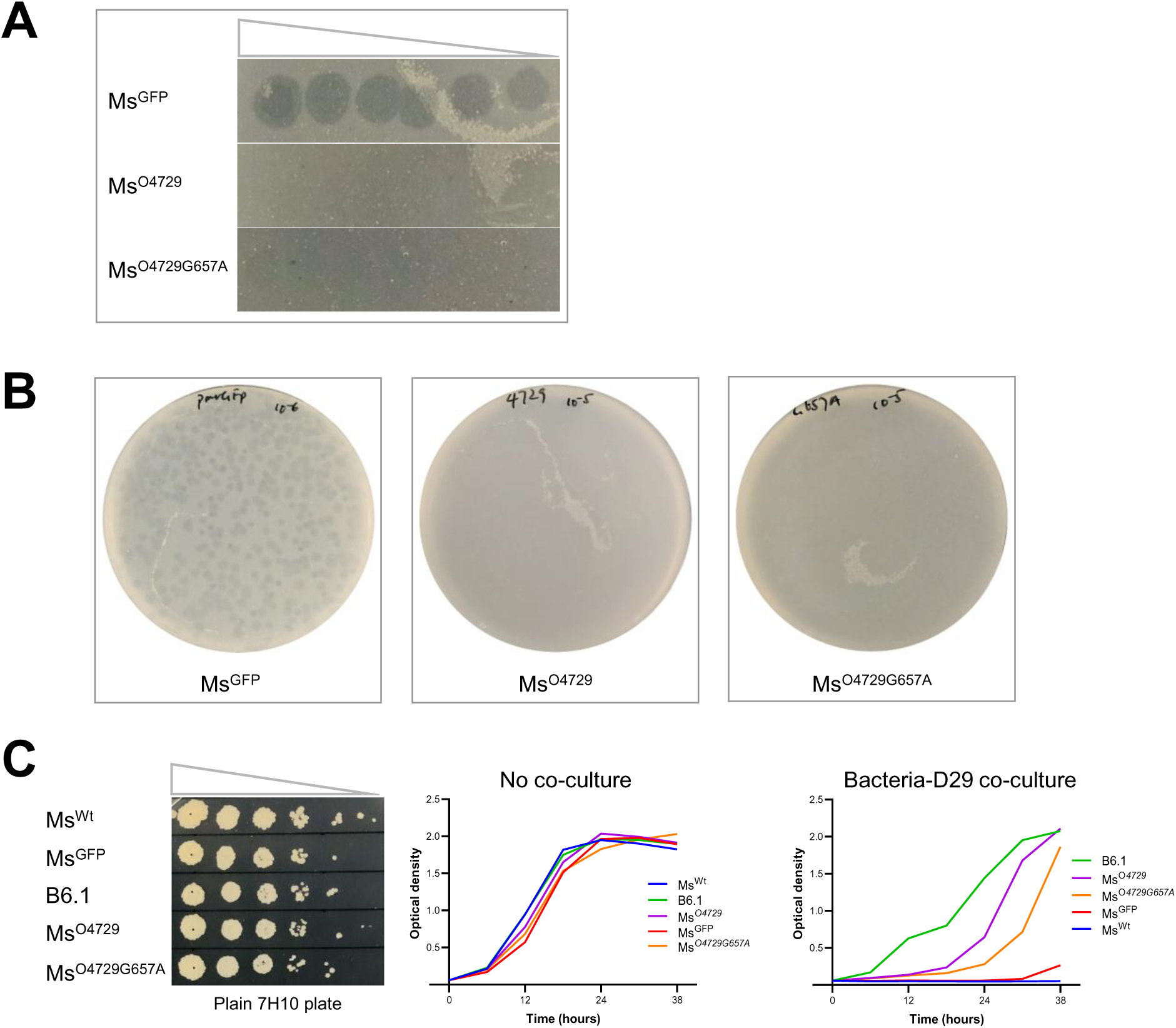
Overexpression of *MSMEG_4729* confers *M. smegmatis* with D29 resistance. A, **Spot-kill assay.** Overexpression of either versions of the gene enables the bacteria to resist formation of zones of growth inhibition during spot-kill assay. B, **Plaque assay.** The overexpression strains also appear to resist plaque formation by D29. C, **Growth curves.** Growth in the absence of D29 infection is similar across all tested strains (left and middle panels) but differs starkly when exposed to D29 infection (right panel). Significant growth is seen in B6.1 and the overexpression strains over a period of 38 hours in the presence of D29 infection.

Given that constitutive overexpression of wild-type *MSMEG_4729* correlated with D29 resistance, we hypothesized that expression levels of this gene or others within the LOS cluster might differ between Ms^Wt^ and B6.1. To test this, we quantified transcript abundance of all LOS cluster genes via RT-qPCR. While baseline expression showed minimal differences between B6.1 and Ms^Wt^ (Figure 12A), D29 exposure triggered widespread up-regulation of the entire LOS cluster in B6.1—most prominently for *MSMEG_4733*, which encodes a putative membrane protein (Figure 12B). This induction aligns with B6.1’s ability to grow under D29 pressure, unlike Ms^Wt^ (Figure 11C), suggesting that phage-responsive transcriptional activation, rather than baseline expression, drives resistance.

**Figure 12.**
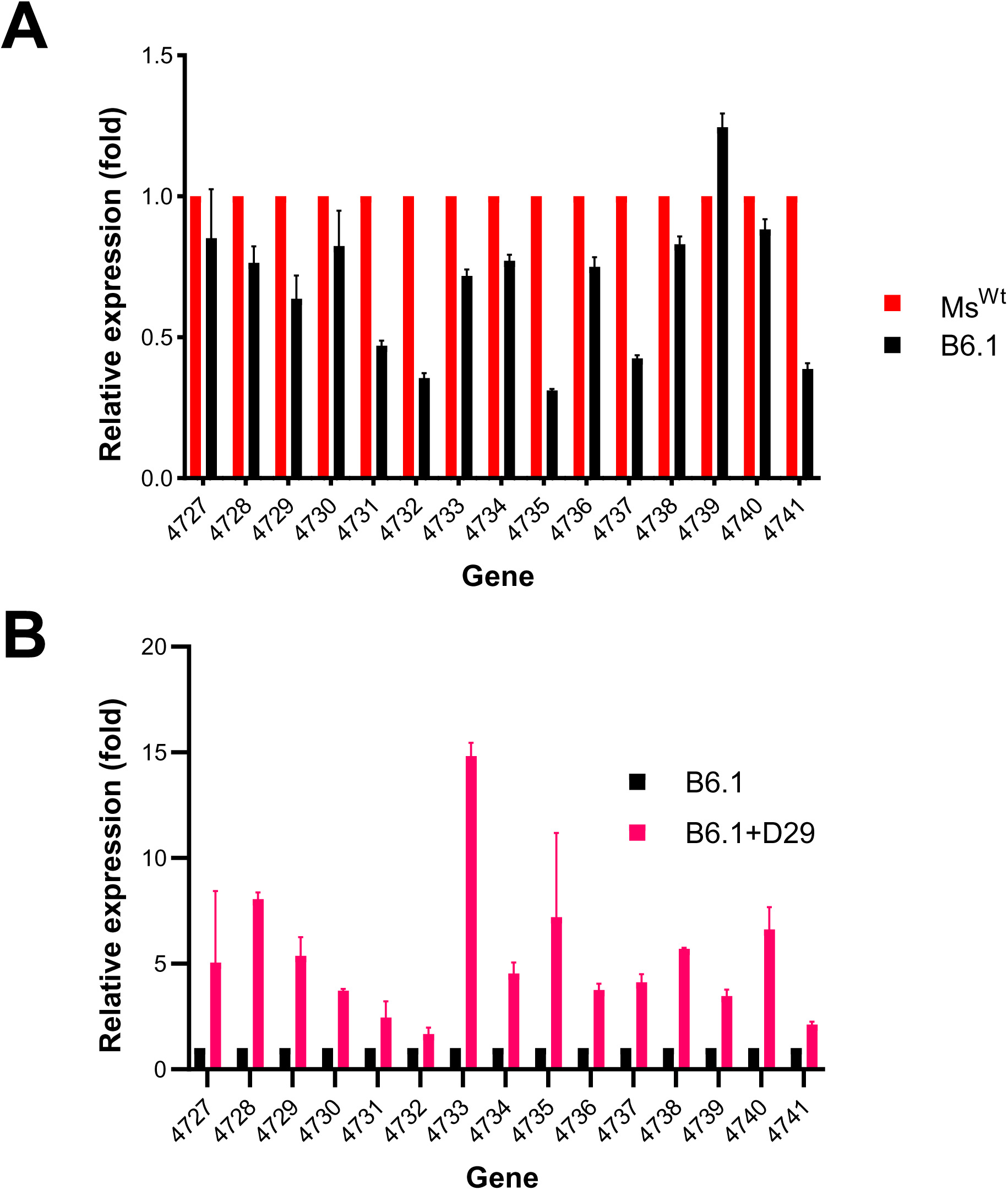
Relationship between D29 resistance and LOS cluster expression profile. A, in the absence of D29 infection, expression of LOS cluster is mostly lower in B6.1 than in Ms^Wt^, except in the case of *MSMEG_4739*. B, LOS cluster is up-regulated in B6.1 following exposure to D29, and expression level is particularly higher for MSMEG_4733, which encodes a putative membrane protein.

### Isolation of DEMs of D29 against B6.1

Wild-type D29 forms turbid zones of growth inhibition on B6.1, and this phenotype persisted during secondary infections using recovered plaques (Figure 7, Figure S5), indicating stable resistance. We generated DEMs via a 28-day co-culture of D29 with Ms^Wt^, and the resulting phage was termed Ms^Wt^-trained <u>D29</u> (tD29). When tD29 was used to infect B6.1, four isolated plaques (tD29-1–4) formed clear zones of inhibition—in stark contrast to the turbid zones formed by wild-type D29 (Figure 13), which suggests that tD29 had evolved enhanced infectivity toward B6.1. Sequencing of tD29-1 and tD29-3 confirmed they were true DEMs, each harboring mutations in multiple genes. Both DEMs shared identical mutations in *gp32* and non-identical changes in *gp14* and *gp30* (Table 3), implicating these loci in overcoming B6.1 resistance.

**Figure 13.**
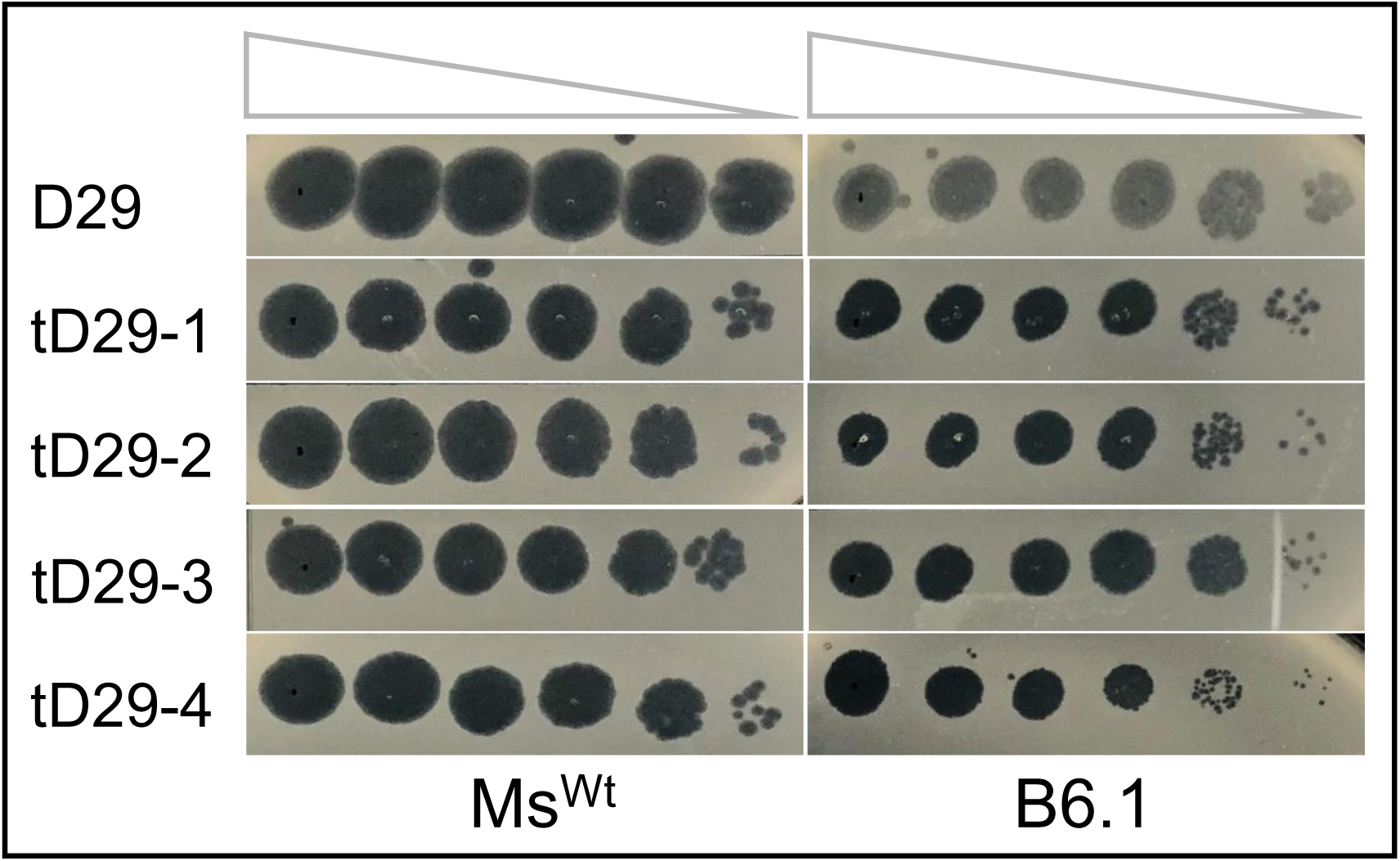
Ms^Wt^-trained D29 (tD29) infection profile on B6.1. D29 forms rather turbid plaques/zones of growth inhibition on B6.1 in contrast to the clearer ones formed on Ms^Wt^ (Figure 7). This is the same for purified plaques recovered from D29 infection of B6.1 (Figure S5). However, D29 from 28-day co-culture with Ms^Wt^ (tD29) was used to infect B6.1, and the plaques were purified and amplified on Ms^Wt^. This purified plaques (tD29-1 through 4) appear to form clear zones of growth inhibition on B6.1 compared to D29 or the purified plaques.

**Table 3.**
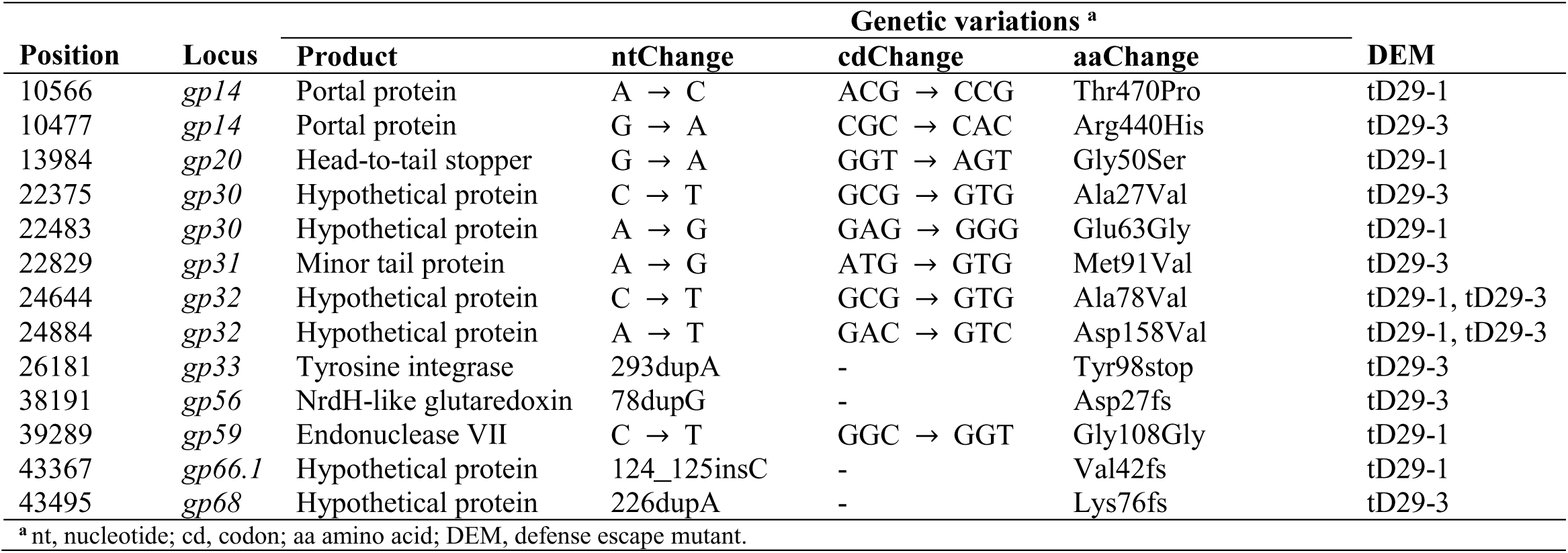
Genetic mutations in DEMs of D29.

### Epigenetic modifications in some D29-resistant mutants

D29 resistance mutations occurred in 18 of the D29-resistant mutants, whereas the remaining 6 had no mutations. However, 3 of the 6 mutants without mutation appear to have sustained IS transposition events (19). In addition, introducing the synonymous mutation *4729G657A* did not confer the bacteria with D29 resistance (Figure 9). We therefore checked for possible epigenetic changes in select strains based on the following criteria: i) the mutant either sustained no mutation or sustained only the synonymous mutation *4729G657A*; ii) the mutant sustained no IS transposition event; and iii) Ms^Wt^ was selected as the control strain. A total of six strains were selected (Table S2).

Using PacBio SMRT sequencing, we detected varying proportions of adenine methylation (m^6^A) within the motif “CTCGAG” across the genomes of all six strains, including Ms^Wt^. The variation in proportions of modified sites (Ms^Wt^, 86.76 %; B6.1, 77.67 %; B11.3, 78.72 %; B15.1, 97.58 %; B54.2, 96.69 %; C77.1.1, 89.77 %) indicates potential loss/gain of m^6^A modifications in these mutants. REBASE search revealed that *M. smegmatis* mc² 155 encodes two restriction-modification (r-m) proteins: MSMEG_3213 (type II methyltransferase that recognizes “CTCGAG”) and MSMEG_3214 (type II restriction endonuclease that recognizes “CTYRAG”) (Table S3–S4) (23). The observed m^6^A profile aligns with MSMEG_3213 activity, likely implicating r-m systems in anti-phage defense. However, the link between the detected m^6^A modification pattern and D29 resistance remains to be uncovered.

### Genome-wide prediction of anti-phage defense systems in *M. smegmatis*

Prokaryotic genomes encode diverse anti-phage defense systems. Using DefenseFinder (24), we identified 7 predicted defense systems in *M. smegmatis*, including the two r-m proteins MSMEG_3213-3214 (Table 4). Notably, only a limited overlap existed between these predictions and the likely defense systems identified in this study. For example, while the Ceres system (MSMEG_5390) was not exactly detected in the spontaneous mutants, its neighboring operon gene *MSMEG_5393* sustained disruption by ISMsm1 transposition in one of the resistant mutants (19). Collectively, these findings reveal that beyond known defense pathways, novel genetic factors may contribute to D29 resistance in *M. smegmatis*.

**Table 4.** Genome-wide prediction of anti-phage defense systems in *M. smegmatis*.

| System | Gene(s) | Product |
| --- | --- | --- |
| Wadjet | *MSMEG_0368* | hypothetical protein |
|  | *MSMEG_0369* | hypothetical protein |
|  | *MSMEG_0370* | conserved hypothetical protein |
| SspBCDE | *MSMEG_1243* | conserved hypothetical protein |
|  | *MSMEG_1244* | conserved hypothetical protein |
|  | *MSMEG_1245* | phosphoadenosine phosphosulfate reductase |
| Prometheus | *MSMEG_1252* | conserved hypothetical protein |
| Wadjet | *MSMEG_1279* | conserved hypothetical protein |
|  | *MSMEG_1280* | hypothetical cytosolic protein |
|  | *MSMEG_1281* | conserved hypothetical protein |
|  | *MSMEG_1282* | hypothetical cytosolic protein |
| Type II RM (Type II MTases) | *MSMEG_3213* | conserved hypothetical protein, Probable DNA methylase |
| Type II RM (Type II REase) | *MSMEG_3214* | hypothetical protein |
| Ceres | *MSMEG_5390* | conserved hypothetical protein |
| Esos | *MSMEG_5849* | conserved hypothetical protein |
| MTase, methylase; REase, restriction endonuclease. |  |  |

## Discussion

Phage therapy constitutes a promising, safe alternative to antibiotic therapy, particularly in the face of rising threat of AMR. Bacteriophages have demonstrated remarkable potential for utility against bacterial infections, including in complicated cases of failed antibiotic therapy (3,25,26). However, to ensure effective, sustainable clinical utility of phages against bacterial infections, certain challenges must be addressed, including phage resistance. Bacterial pathogens have developed a wide range of defensive measures against bacteriophages, which constitute an impediment to phage therapy. Therefore, dissecting the mechanisms of phage-host interactions would be invaluable in uncovering bacterial anti-phage defenses and ways of overcoming them, hence guiding rational phage engineering efforts for therapeutic applications (3).

Transposon mutagenesis is a powerful genome-wide approach to defining genetic factors underlying phage-bacteria interactions (27), but when those genetic factors are essential to bacterial survival, they may often be missed by this approach. Phage-bacteria interaction is greatly influenced by cell envelope-associated mycolic acids, lipids and sugars (12,14,28–33). Genes involved in cell envelope biogenesis as well as lipid metabolism are often essential in mycobacteria, thus limiting the coverage of transposon mutagenesis in defining key aspects of phage-mycobacteria interactions. Here, we report a high frequency of spontaneous D29 resistance in *M. smegmatis*. In a representative sample of 24 D29-resistant strains, we have shown that even if there have been any cell envelope changes, they do not appear to manifest in surface morphology or inhibit D29 adsorption (Figure 2). It could be inferred therefore that despite resistance, surface receptors on the D29-resistant mutants remain accessible to the phage.

Under strong selection, microbes adopt adaptive strategies such as gene duplication or amplification (34–36), but when such mechanisms are not advantageous, they may resort to mutations. We identified mutations in two non-coding regions and 21 genes involved in various pathways in the D29-resistant mutants, including a missense mutation in *MSMEG_4724* - an essential gene that encodes an oligoribunuclease (Table 1). Mutations in the non-coding regions did not introduce new elements to the mutation sites. Some D29-resistant mutants sustained as many as 3-4 mutations, which could have resulted from the multiple exposures to D29. Multiple mutations correlate to greater bacterial fitness in the face of phage pressure, particularly when the mutation introduces a frameshift in the peptide sequence (37). Each of A72.1 and 2.6.1 - the two most resistant mutants (Figure 2) - sustained 3 mutations, with one frameshift in a putative peroxidase and a bacterioferritin comigratory protein respectively (Figure 3, Table 1). However, 5.2.1 - which is not as resistant as A72.1 and 2.6.1 - sustained 4 mutations, including two frameshifts in a glycosyltransferase and the large subunit of exodeoxyribonuclease 7. This suggests that in addition to the mutations and frameshifts in A72.1 and 2.6.1, other factor(s) may be responsible for their remarkably high D29 resistance phenotype (19).

We detected high frequency of *4729G657A* mutation, which is a nucleotide substitution where a codon TTG was changed to TTA in *MSMEG_4729*. Both codons encode a leucine residue, but TTA is a rarely used codon in *M. smegmatis*. While synonymous mutations are often perceived to be innocuous, they could play particularly significant roles in bacterial fitness (38–40). We identified the D29-resistant mutant B6.1 that sustained only this synonymous mutation, then constructed a knockout strain for *MSMEG_4729* and another strain in which the synonymous mutation had been introduced by targeted-base editing. Both knockout and base-edited strains for *MSMEG_4729* have similar D29 susceptibility profiles as Ms^Wt^ (Figure 9), suggesting that deletion of the gene does not cause resistance and the synonymous mutation is not the sole factor responsible for the D29 resistance phenotype.

Effects of synonymous mutations are mostly neutral under normal conditions (41), but may be amplified under strong selection (42). Therefore, we subjected the base-edited strain - alongside B6.1 and the knockout strain as controls - to a 72-hour D29 exposure in liquid medium. While B6.1 grew under this condition, the base-edited and knockout strains did not, which rules out the possibility that the synonymous mutation plays role in D29 resistance in an adaptive fashion. Synonymous mutations could have various effects, which include increasing translation efficiency (42), up-regulation (43) or down-regulation (44) of protein expression, promoter reconstitution and increased mRNA stability (39,45,46), etc. We have seen no evidence of promoter reconstitution or change in secondary mRNA structure at the mutation site. We therefore checked whether overexpression of wild type or mutated *MSMEG_4729* could confer *M. smegmatis* with D29 resistance. *M. smegmatis* became obviously resistant to D29 when either of the genes was overexpressed (Figure 11). This raises the question of whether the D29 resistance phenotype of B6.1 could be associated with overexpression of *MSMEG_4729*. *MSMEG_4729* lies in a 15-gene LOS cluster that is associated with phage resistance in *M. smegmatis* (12,22,47). For this reason, we quantified the expression of not just *MSMEG_4729*, but all 15 genes in both Ms^Wt^ and B6.1 by RT-qPCR. There were mostly minimal differences in transcript levels of these genes in B6.1 compared to those in Ms^Wt^ (Figure 12A). However, following B6.1 exposure to D29, all the genes became up-regulated (Figure 12B), especially *MSMEG_4733*, which encodes a putative membrane protein. This could suggest that expression of the cluster is activated in B6.1 as an adaptive response to D29 exposure, which likely depends on an unknown factor that remains to be uncovered.

The LOS cluster–known for its role in LOS biosynthesis–is negatively regulated by Lsr2 in *M. smegmatis* such that activation of the cluster would require disruption of *lsr2* (12,13). However, not all strains of *M. smegmatis* produce LOS. While *M. smegmatis* ATCC 607 produces this class of lipids (21,22), activation of this cluster in *M. smegmatis* mc^2^ 155 is associated with accumulation of PIMs (12). Exposure to the mycobacteriophage TM4 triggers the transposition and integration of IS1096 into *lsr2*, which disrupts *lsr2* and activates the LOS cluster, resulting in PIMs accumulation and phage-resistant *M. smegmatis* mc^2^ 155 strains with smooth surface morphology (12). We have seen no evidence of mutation or disruption of either *lsr2* or its paralog *MSMEG_1060* (47) in any of the 24 D29-resistant mutants, and it is not clear whether this could explain the lack of change in surface morphology of the D29-resistant mutants in this study.

Moreover, a relationship has been drawn between the LOS cluster, *lsr2*, *MSMEG_4242* and *pknL (MSMEG_4243)* in *M. smegmatis* mc^2^ 155. Disruption of *pknL* and *MSMEG_4242* results in increased expression of the IS1096 transposase TnpA, *lsr2* inactivation, expression of LOS genes, lipid accumulation and smooth surface morphology (13). *MSMEG_4242* and *pknL* are located adjacent to each other, just downstream of a two-gene operon–*MSMEG_4240-4241*. There is a 66 bp non-coding region between *MSMEG_4241* and *MSMEG_4242*, which contains three 19-bp sequences that appear nowhere else on the entire *M. smegmatis* genome. In this study, we identified a D29-resistant mutant with 3 mutations, including *4729G657A* and a 25-bp deletion (including one of the 19-bp sequences) from this region, leaving behind the remaining two 19-bp sequences separated and flanked on both sides by identical 5-nucleotide sequences (Figure 5). The LOS cluster and *MSMEG_4242-pknL* clearly share a connection via *lsr2* inactivation, and both appear to be relevant to mycobacterial lipid metabolism and membrane biogenesis, but it remains to be seen whether they could also be related to the *MSMEG_4240-4241* operon in anyway. *MSMEG_4240* encodes a polyprenyl synthetase, which could likely be involved in the synthesis of polyprenyl phosphates for transmembrane transport of sugar units during membrane biogenesis (48). *MSMEG_4241* and *pknL* possess 14 and 1 transmembrane domains respectively, suggesting possible association with the membrane. *MSMEG_4242* encodes a putative regulatory protein. Therefore, the fact that these four genes are located adjacent to each other, coupled with the occurrence of *4729G657A* and *MSMEG_4241-4242* fusion in the same D29-resistant mutant, could suggest possible coordination in lipid and cell envelope metabolism in *M. smegmatis*. However, this remains to be validated by further experimentation.

We mentioned earlier that *M. smegmatis* mc^2^ 155 does not produce LOS, but its parent strain–*M. smegmatis* ATCC 607–does. However, the LOS clusters in both strains share a 99.99 % similarity at nucleotide level. The mc^2^ 155 LOS cluster appears to have two “C” insertions and a transmembrane domain in the N-terminal of MSMEG_4734 (Figure 8B). If the N-terminal of *NCTC7017_04217*–the ATCC 607 ortholog of *MSMEG_4734*–were to be extended by 186 bp (assuming no sequencing error and that the difference in protein length is due to possible misannotation), the new start position (CGA) for the gene wouldn’t correspond to a start codon. In addition, if the N-terminal were to be extended by 184 bp (taking into account the 2 gaps introduced by the two “C” insertions uncovered by the sequence alignment), the new start position (ATG) corresponds to a start codon, but 184 bp is not a multiple of 3. Even if we were to assume another possible alternative start position within the upstream region of *NCTC7017_04217*, the only such possible position would be at -108 bp. This new sequence would therefore translate into a protein that wouldn’t include the entire 22-amino acids transmembrane domain predicted in MSMEG_4734. Thus, it appears that the only difference between the two clusters is the two insertions that seem to be later acquired by the mc^2^ 155 strain, which is significant enough to introduce an N-terminal transmembrane domain in the peptide sequence of MSMEG_4734. How this could be part of the reason for lack of LOS synthesis by the mc^2^ 155 strain remains to be experimentally checked.

Phage malleability is one of the advantages of phage therapy over conventional antibiotic therapy. Unlike drugs–which become ineffective once the bacteria develop resistance to them–phages can mutate to keep up with corresponding changes in the host bacteria, hence maintaining their capacity to infect and kill the host provided the resistance mechanism could be overcome. This process is called defense escape. Though B6.1 clearly manifests high level of resistance to D29, it allows for formation of plaques, albeit to a far lesser extent than Ms^Wt^ (Figure 7A). These plaques, when purified, show identical infection profile as wild type D29 on B6.1 (Figure S5), suggesting that they are more likely to have appeared due to leaky resistance than defense escape (49). However, co-evolutionary phage training of D29 on Ms^Wt^ yielded DEMs of D29 that infect B6.1 with far greater efficiency than the wild type phage (Figure 13), suggesting that the mechanism of D29 resistance in B6.1 could be overcome. We sequenced two DEMs tD29-1 and tD29-3, and of all detected mutations, only *gp32* mutations are identical between both DEMs (Table 3). A non-identical mutation in *gp32* is proposed to mediate D29 escape from vector-borne Mpr-overexpressing *M. smegmatis*, suggesting that Gp32 could be particularly relevant to D29-*M. smegmatis* interactions (18). Further studies will consider dissecting the B6.1 resistance mechanism, how the phage mutations mediate D29 escape, and possible impacts of the mutations on D29 itself.

Some of the sequenced strains harbor neither mutations nor IS transposition events despite D29 resistance (19). Some have the *4729G657A* synonymous mutation only, which we determined is not the sole factor responsible for D29 resistance. It is evident in Mpr-driven D29 resistance that phage resistance may arise in the absence of any mutation (16,17). It was always believed that there had to be a genetic factor behind Mpr overexpression. Likewise, there has to be an unknown factor behind D29 resistance in those strains harboring only the synonymous mutation or no changes at all. Our data suggests that this factor may have nothing to do with either point mutation or IS transposition. However, r-m systems are quite prevalent anti-phage defense systems in bacteria (50), so we checked for possible activity of these systems in the strains. We identified the likely activity of a known type II methyltransferase (MSMEG_3213) in all strains including Ms^Wt^ by the presence of m^6^A modifications. Computational prediction of *M. smegmatis*-encoded anti-phage defense systems also uncovered this methyltransferase and its cognate type II restriction endonuclease (MSMEG_3214) (Tables 4 and S4). It appears that B15.1, B54.2 and C77.1.1 acquired more m^6^A modifications in addition to those already in the wild type strain. Though B6.1 and B11.3 appear to have less proportion of modified sites, loss/gain of m^6^A modifications is possible. Further work will be required to investigate the role of these modifications in *M. smegmatis* resistance to D29.

Growth curves in liquid medium suggest minimal differences between Ms^Wt^ and B6.1 in the absence of D29 infection (Figure 11C, middle panel), which may suggest that the D29 resistance phenotype exerts no fitness effect on B6.1. However, estimation of maximum specific growth rate (µ_max_) per hour (h^-1^) using modified Gompertz equation (51) reveals differences between both strains under both conditions: No co-culture (Ms^Wt^, 0.32 h^-1^; B6.1, 0.21 h^-1^); Bacteria-D29 co-culture (Ms^Wt^, 1.86E-14 h^-1^; B6.1, 0.08 h^-1^). The growth rates in the absence of co-culture therefore indicate possible fitness effect associated with B6.1 D29 resistance phenotype.

This study exposed a wide range of genetic factors that may be involved in the interaction between *M. smegmatis* and the lytic mycobacteriophage D29. We report that exposure to D29 triggers point mutations and epigenetic modifications across the genome. A synonymous mutation (Leu219Leu) occurs at a high frequency, and a strain with this mutation seems to demonstrate Lsr2-independent activation of LOS cluster. Further inquiries will be required to uncover the mechanisms and roles of both *M. smegmatis* and D29 mutations in their interactions, which will inform phage engineering efforts towards building a robust pipeline of therapeutic mycobacteriophages.

## Materials and Methods

### Phage, bacterial strains and growth conditions

The lytic mycobacteriophage D29 was used for all phage experiments in this study, and all infection experiments were carried out at 37 °C in the presence of 2 mM CaCl_2_. *E. coli* Trans1 was used for all cloning experiments, and was maintained on solid or in liquid Luria-Bertani medium in the presence of selective antibiotics where necessary (kanamycin 50 µg/mL, ampicillin 100 µg/mL or zeocin 30 µg/mL). *M. smegmatis* mc^2^ 155 cells were maintained in Middlebrook 7H9 medium or on 7H10 agar. Middlebrook 7H9 medium was supplemented with 10% OADC, 0.2 % glycerol and 0.05% Tween 80, whereas the 7H10 was supplemented with 10% OADC and 0.5 % glycerol. Selective antibiotics were added where necessary (kanamycin 20 µg/mL (liquid) or 50 µg/mL(solid), zeocin 30 µg/mL). All liquid cultures were maintained at 37 °C with shaking at 200 revolutions per minute (rpm).

### Phage amplification

We have previously described the phage amplification method (19). Briefly, exponential phase wild type *M. smegmatis* cells (Ms^Wt^) were pelleted and washed three times in MP buffer, then re-suspended in the same buffer. A mixture of 10-fold serially diluted D29, washed Ms^Wt^ cells, 2 mM CaCl_2_ and top agar was prepared, dispensed into plain 7H10 plates and spread evenly. The preparation was allowed to dry, followed by incubation at 37 °C for 12-24 hours. Amplified phage was recovered in MP buffer, filtered through a 0.22 µm filter and stored at 4 °C until use.

### Generation of spontaneous D29-resistant strains

Spontaneous D29-resistant strains were generated as described previously (19,27,52). Briefly, the strains were generated via multiple rounds of D29 exposure at an MOI of 5, with the exposure time extended by 7 hours after each round. The infection medium used was MP buffer, and after each exposure cycle, the cells were washed in PBS-Tween 80, sub-cultured in 7H9 medium containing Tween 80 until exponential phase, then washed in MP buffer and subjected to the next round of infection. Cells were washed in PBS-Tween 80 after the final round of exposure, then serially diluted, streaked on plain 7H10 plates and incubated at 37 °C until the appearance of single colonies.

### Phage susceptibility testing (PST)

Susceptibility of bacterial strains to D29 was evaluated by plaque assay, spot kill assay and their ability to grow on D29-seeded 7H10 plates as was described previously (19). Briefly, plaque assay was prepared by pouring a soft overlay of top agar, D29, MP buffer-washed Ms^Wt^ cells and 2 mM CaCl_2_ on three plain 7H10 plates, then allowed to dry prior to incubation at 37 °C for 12-24 hours before enumeration of plaque-forming units (PFU). Spot kill assay was first prepared by pouring a soft overlay of top agar, MP buffer-washed Ms^Wt^ cells and 2 mM CaCl_2_, then allowed to dry. This was followed by spotting of ten-fold serial dilutions on D29 on the dried plate, allowed to dry, and incubated at 37 °C for 12-24 hours. For PST on phage-seeded plates, ten-fold serial dilutions of washed cells were prepared in MP buffer containing 2 mM CaCl_2_, 2 µL of which were spotted on a 7H10 plate (pre-seeded with 10^7^ – 10^9^ PFU/milliliter of D29). The plate was dried and incubated at 37 °C for 3-5 days. A negative control was prepared by plating serial dilutions of the washed cells on a plain 7H10 plate.

### Phage adsorption assay

Phage adsorption assay was performed according to previously described protocols (12,27). Mid-log phase bacterial cells were washed and re-suspended in MP buffer, then diluted to OD_600_ 0.3. Terminal adsorption was determined by adding 30 µL (10^-3^ dilution) of D29 to 200 µL of diluted bacteria, followed by addition of 2 mM CaCl_2_. A similar preparation was simultaneously prepared by adding the same volume of phage at the same dilution to 200 µL of sterile MP buffer, followed by addition of 2 mM CaCl_2_. The first preparation was incubated at 37 °C with shaking for 60 minutes before collecting the supernatant for subsequent dilution and plaque assay to determine the percentage of adsorbed phage, whereas the supernatant was immediately collected from the second preparation to enumerate the input PFU. Similar preparations were prepared for enumeration of PFU at 0-, 30- and 60-minute time points to determine the percentage of unadsorbed phage (presented as phage adsorption rate).

### Whole genome sequencing (WGS) and data analysis

Genomic DNA (gDNA) extraction and subsequent sequencing were performed by Shanghai Jingnuo Biotechnology Co., Ltd. (Shanghai, China). gDNA was isolated using a modified cetyltrimethylammonium bromide (CTAB) protocol (53). The extracted DNA was quantified with a Qubit 2.0 Fluorometer (ThermoFisher, USA). Library preparation was carried out using the Hieff NGS^®^ OnePot Flash DNA Library Prep Kit (Yeasen, China), which involved enzymatic fragmentation, end repair, A-tailing, adapter ligation, purification, and PCR amplification. Following an initial quantification with the Qubit 2.0 and appropriate dilution, fragment size distribution was assessed on an Agilent 2100 Bioanalyzer (Agilent, USA). The effective concentration of each library was then precisely determined by qPCR. Libraries meeting quality standards were pooled, clustered on a cBOT instrument, and sequenced on the NovaSeq 6000 platform (Illumina, Inc., USA).

Next generation sequencing data generated from WGS was analyzed using the GalaxyEU bioinformatics platform, with settings set to default parameters (54). Quality of raw sequence reads was checked using FASTQC (55), and trimming was carried out using FASTP (56). Snippy (57) was used to map trimmed reads to the reference genome (NCBI accession number CP000480.1) and call variants. Sanger sequencing was used to verify all detected variants.

### Genome editing

Genome editing was performed by CRISPR/Cas12a-assisted recombineering via the homology-directed repair (HDR) pathway (58). The plasmid pJV53-Cpf1 was first transformed into *M. smegmatis* to obtain Ms:pJV53-Cpf1. CRISPR RNAs (crRNAs) were designed from *MSMEG_4729* by identifying 23 bp sequences (with a GC content of 40-60%) downstream of the protospacer adjacent motif “YTN” (59). *Bpm*I and *Hin*dIII sticky ends were added to the 23 bp crRNA sequence, which was labeled cr*4729*<u>k</u> (for <u>k</u>nockout) or cr*4729*<u>e</u> (for targeted base <u>e</u>diting) and synthesized as forward and reverse oligonucleotides. The synthesized oligonucleotides for cr*4729*k as well as cr*4729*e were first annealed to obtain double-stranded cr*4729*k and cr*4729*e, then cloned into *Bpm*I/*Hin*dIII-linearized pCR-Zeo to obtain pCR*4729*k and pCR*4729*e respectively.

To construct the homologous repair template (HRT) for knockout, <u>u</u>pstream (U, 800 bp) and <u>d</u>ownstream (D, 827 bp) sequences were amplified at opposite sides of cr*4729*k, including flanking sequences from the adjacent genes *MSMEG_4728* and *MSMEG_4730*. Both arms were cloned into *Eco*RI/*Bam*HI-linearized pBlueSK to obtain the vector pBlueSK-*4729UD* in which the U and D arms are fused into a single double-stranded HRT. The HRT (*4729UD*) was amplified from pBlueSK-*4729UD*, purified and kept for downstream applications. For the targeted base editing, a 59 bp <u>s</u>ingle-<u>s</u>tranded <u>edi</u>ting template for *MSMEG_4729* (ss*4729*ediT) was designed from the plus strand of target gene in which the target nucleotide “G” was replaced by “A”, and “A” is flanked by two 29 bp sequences at both sides. The 23 bp sequence of cr*4729*e was designed from the minus strand and is entirely complementary to ss*4729*ediT.

For the knockout, 2-4 µg each of pCR*4729*k and *4729*UD were transformed into electrocompetent Ms:pJV53-Cpf1 cells by electroporation. Transformed cells were transferred to 2 mL of fresh 7H9 medium and incubated at 37 °C for 5 hours with shaking, then concentrated and plated on 7H11 plates containing kanamycin (50 µg/mL), zeocin (30 µg/mL) and anhydrotetracycline (50 ng/mL). Plates were incubated at 30 °C for 4-5 days, and colonies were picked and verified by PCR and Sanger sequencing. For targeted base editing, pCR*4729*e and ss*4729*ediT were transformed into electrocompetent Ms:pJV53-Cpf1 cells as crRNA and HRT respectively. Base editing was verified by Sanger sequencing.

### Growth kinetics

For growth in the absence of D29, strains were grown in 7H9 medium to OD_600_ 0.8-1 at 37 °C with shaking, then diluted to OD_600_ 0.06 in the same medium. For growth in the presence of D29 infection, all strains were grown in 7H9 medium to OD_600_ 0.8-1 at 37 °C with shaking, then pelleted and washed thrice in 7H9 medium without Tween 80. The washed cells were diluted to OD_600_ 0.06 in 7H9 medium (without Tween 80) containing 2 mM CaCl_2_ and D29 at MOI of 5. The cells were incubated with shaking at 37 °C, and optical density of each strain was checked at 6-hour intervals until exponential phase, and the readings were plotted as growth curves. Simultaneously, ten-fold serial dilutions of non-infected cells were prepared in 7H9 medium, and 2 µL of each were spotted on a plain 7H10 plate, allowed to dry and incubated at 37 °C for 3-5 days before observation.

### RNA extraction and RT-qPCR

RNA extraction, reverse transcription and RT-qPCR were carried out using commercially available kits according to manufacturer instructions. RNA was extracted using Magen HiPure Bacterial RNA Mini Kit (Magen, China), reverse transcription was performed using ExonScript All-in-One RT SuperMix with DNase (EXONGEN, China), whereas RT-qPCR was carried out using Taq Pro Universal SYBR qPCR Master Mix (Vazyme Biotech., China). For quantification of mRNA transcripts in the absence/presence of D29 infection: the cells were first cultured in 7H9 medium containing Tween 80, washed three times in 7H9 medium without Tween 80, then sub-cultured in 7H9 containing 2 mM CaCl_2_ but no Tween 80. D29 was added at an MOI of 5 where needed. All qPCR reactions were run in C1000 Touch^TM^ Thermal Cycler (Bio-Rad, USA). The housekeeping gene *sigA* served as an internal control for normalization. Relative gene expression in the wild-type/untreated control groups was calibrated to a baseline value of 1. Expression levels in the treated/test groups were subsequently calculated as fold change relative to this baseline. All data were visualized as bar graphs using GraphPad Prism version 9.0.0 (GraphPad, USA).

### Phage-bacteria co-evolution experiment

Ms^Wt^ was grown in 7H9 medium to OD_600_ 0.8-1 at 37 °C with shaking, pelleted and washed thrice in 7H9 medium without Tween 80. The washed cells were re-suspended in fresh 7H9 medium without Tween 80, CaCl_2_ was added to a final concentration of 2 mM, D29 was added at MOI of 7, and the culture was incubated at 37 °C with shaking for 28 days. The culture was centrifuged, the supernatant was collected and filtered through a 0.22 µm filter, labeled Ms^Wt^-trained D29 (tD29) and used as phage stock for plaque assay on B6.1. Four single plaques that appeared on B6.1 were collected, amplified on Ms^Wt^ and designated tD29-1, tD29-2, tD29-3 and tD29-4.

### Phage sequencing

D29, tD29-1 and tD29-3 lysates were concentrated by high speed centrifugation at 26,000 rpm for 1 hour using Optima L-100XP Ultracentrifuge (Beckman Coulter, USA). The supernatant was discarded, and the residual phage was re-suspended in 2 mL of SM buffer. DNA was extracted from the concentrated phage lysates and sequenced on Illumina PE150 NovaSeq platform by Azenta Life Sciences (Suzhou, China).

### Epigenome sequencing

Epigenome sequencing was carried out by Azenta Life Sciences (Suzhou, China). DNA was extracted from exponential phase culture of each strain using Magen HiPure Soil DNA Kit (Magen, China), and library preparation was carried out using SMRTbell^®^ prep kit 3.0 (PacBio, USA), both according to manufacturer instructions. PacBio Sequel IIe platform was used for whole genome sequencing, and SMRT Link software was used to perform base modification detection and motif analysis.

## Acknowledgments

This study was supported by the National Key R&D Program of China (2021YFA1300900), the Major Project of Guangzhou National Laboratory (GZNL2024A01009, GZNL2025C01003), the National Natural Science Foundation of China (82502762).

## Conflict of interest

The authors declare no conflict of interest.

## Data availability

The Illumina sequencing data underlying this article are available in the NCBI Sequence Read Archive at https://www.ncbi.nlm.nih.gov/bioproject/PRJNA1423624. Alternatively, the datasets are also available here: https://www.scidb.cn/en/detail?dataSetId=47ad6fcd07f64adea3cc5a17e246d766.

## Notes

https://www.ncbi.nlm.nih.gov/bioproject/PRJNA1423624

https://www.scidb.cn/en/detail?dataSetId=47ad6fcd07f64adea3cc5a17e246d766

### Competing Interest Statement

The authors have declared no competing interest.

