## Supplemental files for "A synonymous mutation in *MSMEG_4729* occurs at a high frequency in spontaneous D29-resistant mutants of *Mycobacterium smegmatis*"

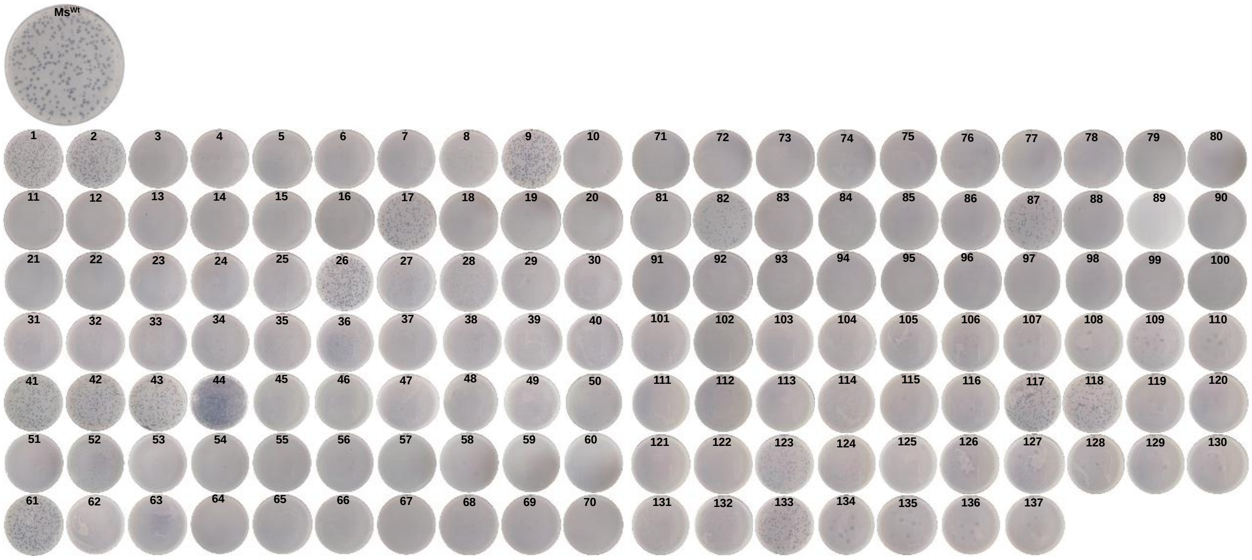


**Figure S1 D29 susceptibility screening.** After four sequential rounds of extended exposures to D29, 137 colonies were picked, grown and screened for D29 resistance. Of these colonies, 21 (15. 3%) maintained their susceptibility, 3 (2.2 %) were partially resistant and 113 (82.5 %) developed phenotypic resistance to D29. Twenty-four (24) of the 137 strains were selected for further purification, susceptibility testing and whole-genome sequencing.

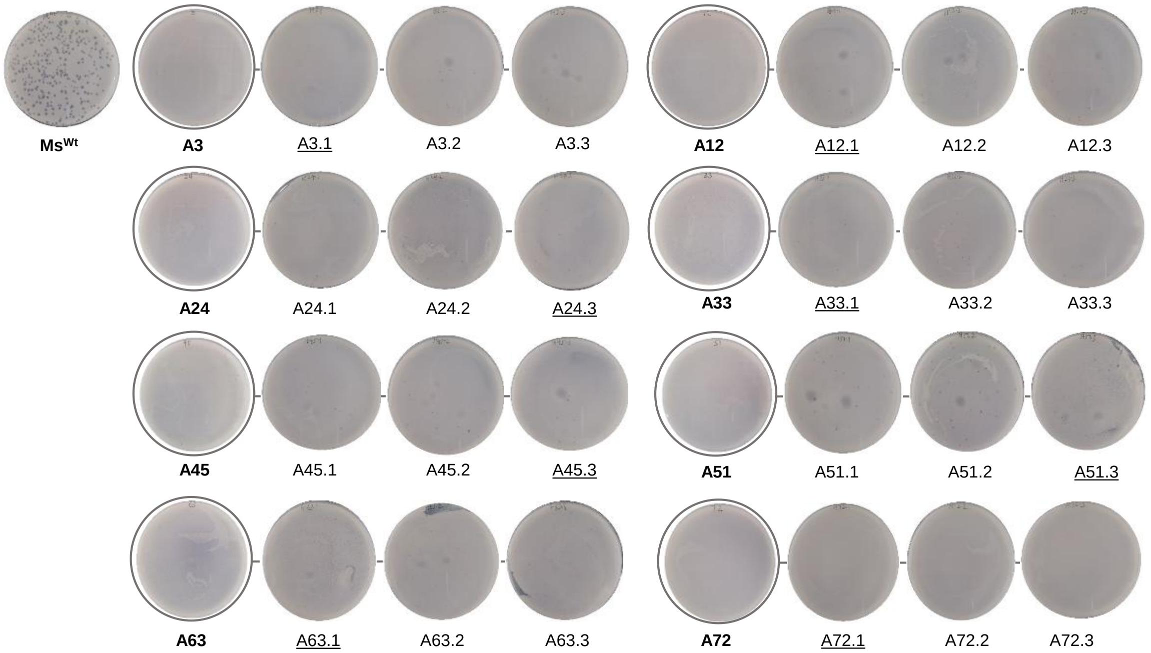


**Figure S2-1 Purification of $1^{st}$ batch of colonies for PST and WGS.** Individual colonies from Figure S1 were screened in three batches A, B and C. This is the first batch (batch A) of screening, and the underlined strains were selected for WGS. PST, phage susceptibility testing; WGS, whole-genome sequencing; A, batch A; 3, colony #3 on Figure S1 (above) and so on; A3.1, colony #1 after first purification of colony number 3 in batch A. The same description and nomenclature applies to Figures S2-2 and S2-3 below, except that the "two periods" in batch C nomenclature means that the strains were purified twice.

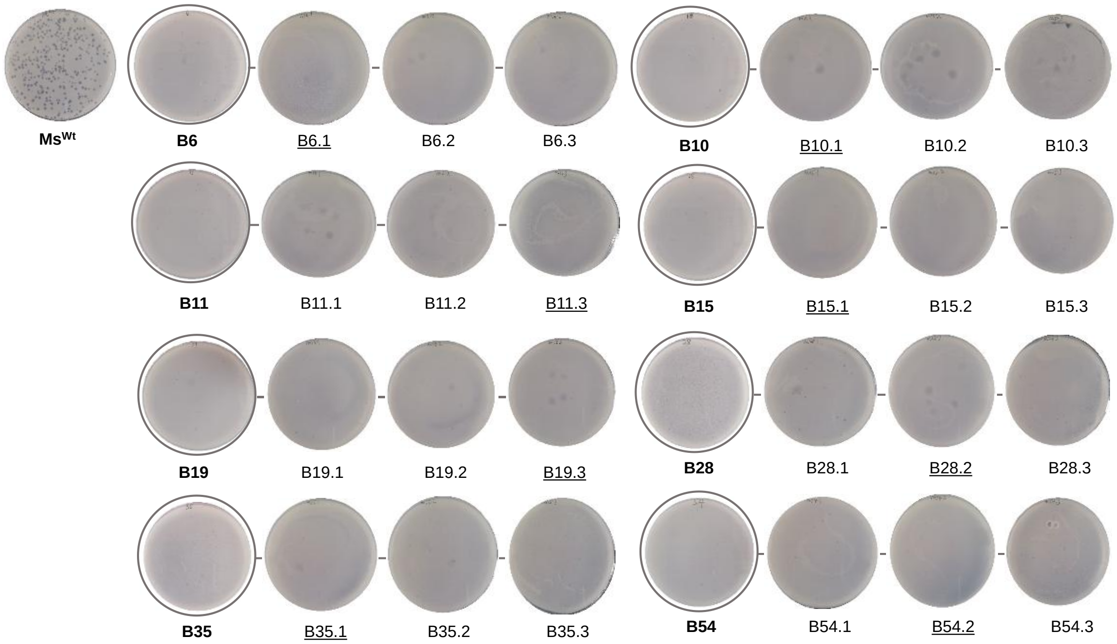


**Figure S2-2 Purification of 2nd batch of colonies for PST and WGS.** See Figure S2-1 above.

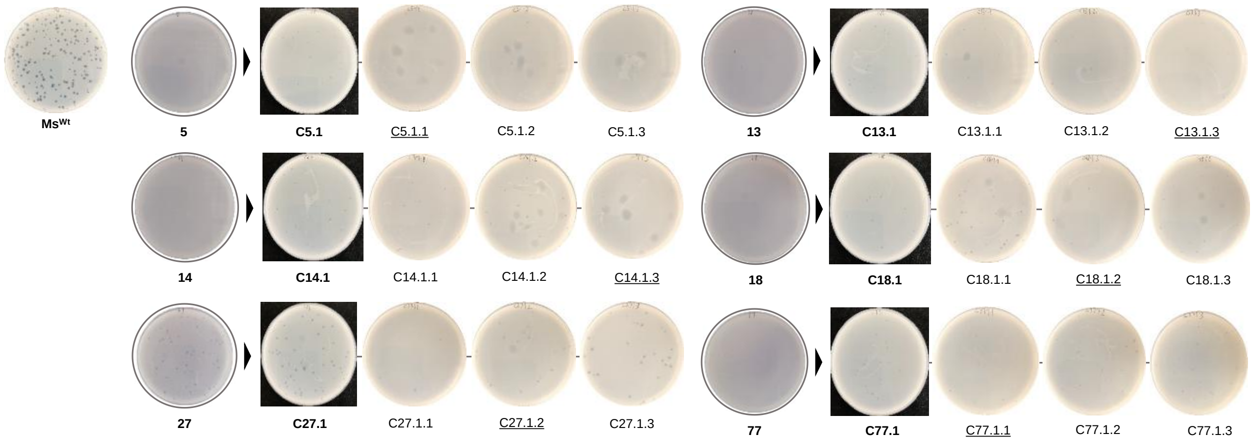


**Figure S2-3 Purification of 3rd batch of colonies for PST and WGS.** See Figure S2-1 above.

**Table S1 Affected gene clusters in the mutant pool**

| S/N | Genes in the cluster [a] | Functional category | Product |
|---|---|---|---|
| 1 | *MSMEG_0016* | [S] | Conserved domain protein |
| | *MSMEG_0017* | [V] | ABC-type multidrug transport system, permease/ATPase |
| | *MSMEG_0018* | [V] | ABC-type multidrug transport system, permease/ATPase |
| | ***MSMEG_0019*** | **[Q]** | **Non-ribosomal peptide synthase (Amino acid adenylaion)** |
| 2 | *MSMEG_0159* | [C] | Formate dehydrogenase, gamma subunit |
| | ***MSMEG_0160*** | **[C]** | **Formate dehydrogenase, beta subunit** |
| | *MSMEG_0161* | [R] | Formate dehydrogenase, alpha subunit |
| | *MSMEG_0162* | - | NAD-dependent formate dehydrogenase, delta subunit |
| | ***MSMEG_0279*** | **[E]** | **Amino acid transporters** |
| | *MSMEG_0278* | - | Hypothetical protein |
| 3 | *MSMEG_0603/fadE6* | [I] | Putative acyl-CoA dehydrogenase |
| | *MSMEG_0604* | [CHR] | Glyoxylate reductase |
| | *MSMEG_0605* | - | Hypothetical protein |
| | ***MSMEG_0606*** | **[K]** | **Transcriptional regulator, TetR family protein** |
| 4 | *MSMEG_0932* | [KG] | Transcriptional regulator/sugar kinase |
| | ***MSMEG_0933*** | **[M]** | **Glycosyltransferase** |
| | *MSMEG_0934* | - | Hypothetical protein |
| | *MSMEG_0935/gpmA* | [G] | Phosphoglycerate mutase 1 |
| 5 | *MSMEG_1279* | - | Hypothetical protein |
| | *MSMEG_1280* | - | Putative cytoplasmic protein |
| | ***MSMEG_1281*** | **[S]** | **Putative cytosolic protein** |
| | *MSMEG_1282* | [S] | Putative cytoplasmic protein |
| | *MSMEG_1283* | [S] | Ribbon-helix-helix transcription factor, family protein |
| | *MSMEG_1284* | [S] | PilT domain-containing protein |
| 6 | *MSMEG_1370* | [ER] | Threonine dehydrogenase and related Zn-dependent dehydrogenases |
| | *MSMEG_1371* | [G] | Phosphomannose isomerase |
| | ***MSMEG_1372*** | **[G]** | **ABC-type sugar transport system, ATPase component** |
| | *MSMEG_1373* | [R] | Predicted ABC-type sugar transport system, permease component |
| | *MSMEG_1374* | [G] | ABC-type sugar transport system, periplasmic component |
| 7 | ***MSMEG_1945*** | **[P]** | **Transmembrane cation transporter** |
| | *MSMEG_1946* | [L] | NADH pyrophosphatase |
| 8 | ***MSMEG_2808*** | **[R]** | **Short-chain dehydrogenase/reductase SDR** |
| | *MSMEG_2809* | - | Hypothetical protein |
| | *MSMEG_2810* | [G] | Arabinose efflux permease |
| 9 | *MSMEG_3764* | [S] | Hypothetical protein |
| | ***MSMEG_3766*** | **-** | **Hypothetical protein** |
| | *MSMEG_4240/idsA2* | [H] | Geranylgeranyl pyrophosphate synthetase/polyprenyl synthetase |
| | ***MSMEG_4241*** | **-** | **Putative integral membrane protein** |
| | *MSMEG_4242* | - | Conserved regulatory protein |
| 10 | ***MSMEG_4724*** | **[A]** | **Oligoribonuclease (3'->5' exoribonuclease)** |
| | *MSMEG_4725* | - | tRNA-His |
| 11 | *MSMEG_4727* | [Q] | Mycocerosic acid synthase |
| | *MSMEG_4728* | - | Condensation domain-containing protein |
| | ***MSMEG_4729*** | **-** | **Hypothetical protein** |
| | *MSMEG_4730* | - | Hypothetical protein |
| | *MSMEG_4731* | [IQ] | Acyl-CoA synthetase (AMP-forming) |
| | *MSMEG_4732* | [R] | Putative glycosyltransferase, group 2 family protein |

| | | | |
|---|---|---|---|
| | *MSMEG_4733* | - | Putative membrane protein |
| | *MSMEG_4734* | - | Hypothetical protein |
| | *MSMEG_4735* | - | Hypothetical protein |
| | *MSMEG_4736* | [S] | Hypothetical protein |
| | *MSMEG_4737* | [S] | Hypothetical protein |
| | *MSMEG_4738* | - | Hypothetical protein |
| | *MSMEG_4739* | - | Hypothetical protein |
| | *MSMEG_4740* | [GC] | Glycosyltransferase family protein |
| | *MSMEG_4741* | [R] | MmpL protein |
| 12 | ***MSMEG_5209*** | **[R]** | **Predicted hydrolases or acyltransferases (alpha/beta hydrolase)** |
| | *MSMEG_5211* | [E] | 4-aminobutyrate aminotransferase and related aminotransferases |
| 13 | *MSMEG_5225* | - | Hypothetical protein |
| | ***MSMEG_5226/xseA*** | **[L]** | **Exodeoxyribunuclease VII, large subunit** |
| | *MSMEG_5227/xseB* | [L] | Exodeoxyribunuclease VII, small subunit |
| 14 | *MSMEG_6001/echA20* | [I] | Enoyl-CoA hydratase |
| | *MSMEG_6002* | [I] | Coenzyme A transferase, alpha subunit |
| | *MSMEG_6003* | [I] | Coenzyme A transferase, beta subunit |
| | *MSMEG_6004* | [R] | Oxidoreductase, 2-nitropropane dioxygenase family protein |
| | ***MSMEG_6005*** | **-** | **Hypothetical protein** |
| | *MSMEG_6006* | - | Hypothetical protein |

[a] Genes that sustained mutations in the D29-resistant mutants are shown in bold font.

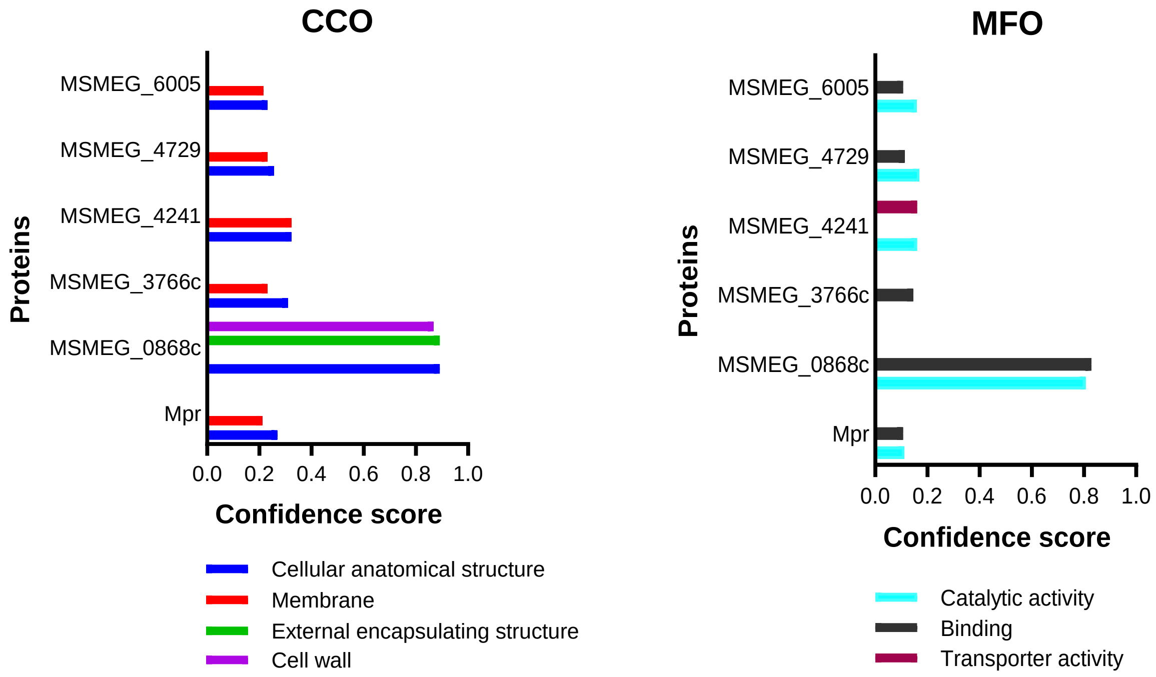


**Figure S3 CCO and MFO prediction for the mutated hypothetical protein-encoding genes.** Five of the mutated genes in the D29-resistant mutants are enriched as hypothetical protein-encoding genes. We carried out function prediction for all of these hypothetical proteins using Mpr as a bioinformatics control. Cellular component ontology (CCO, left) seems to reveal that the proteins may likely be associated with the cell wall/membrane. Molecular function ontology (MFO, right) seems to suggest that all the proteins (except MSMEG_3766c) may have catalytic functions, and MSMEG_4241 may likely be associated with transmembrane transport. Protein functions were predicted using DeepGOWeb (1).

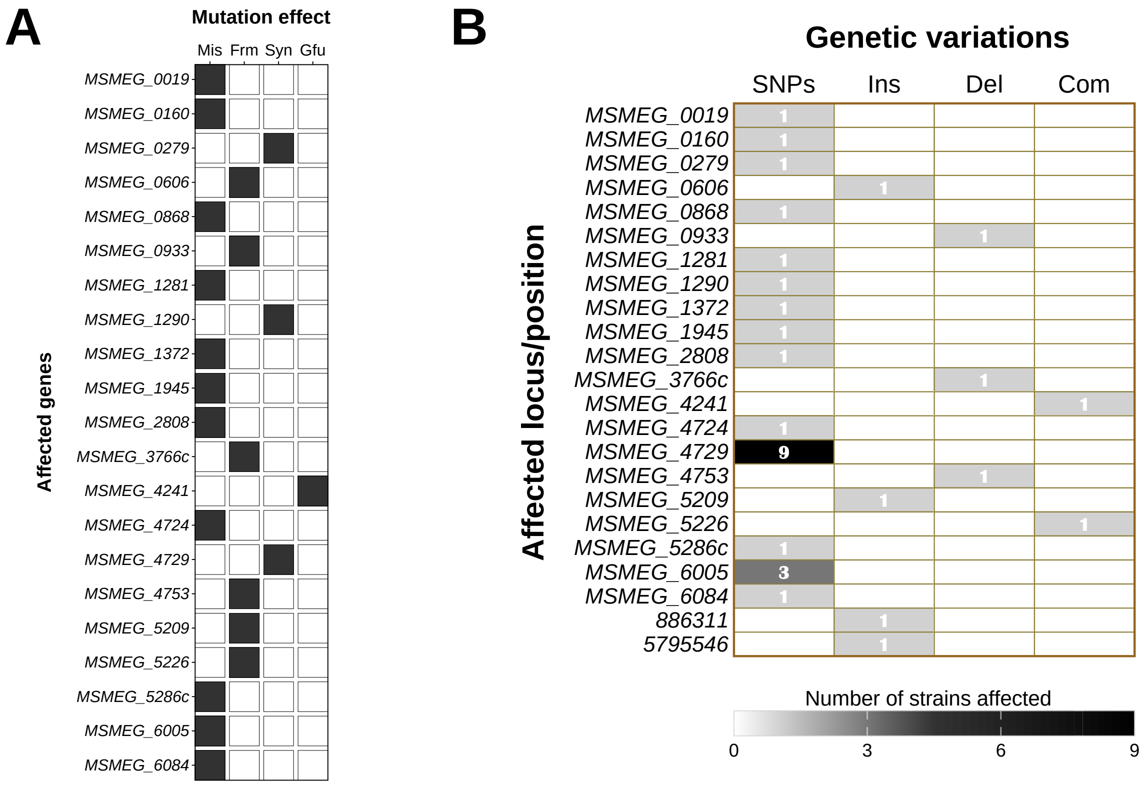


**Figure S4 Mutated genes, mutation effects/types and frequency of occurrence.** Varying effects (Mis, Frm, Syn and Gfu) (A) arose across genomes of the different D29-resistant mutants due to different kinds of mutations (SNPs, Ins, Del, Com) (B). A nucleotide substitution with synonymous effect on MSMEG_4729 had the highest frequency of occurrence (9/24), followed by MSMEG_6005 (3/24). Mis, missense; Frm, frameshift; Syn, synonymous; Gfu, gene fusion; SNPs, single nucleotide polymorphisms; Ins, insertion; Del, deletion; Com, complex.

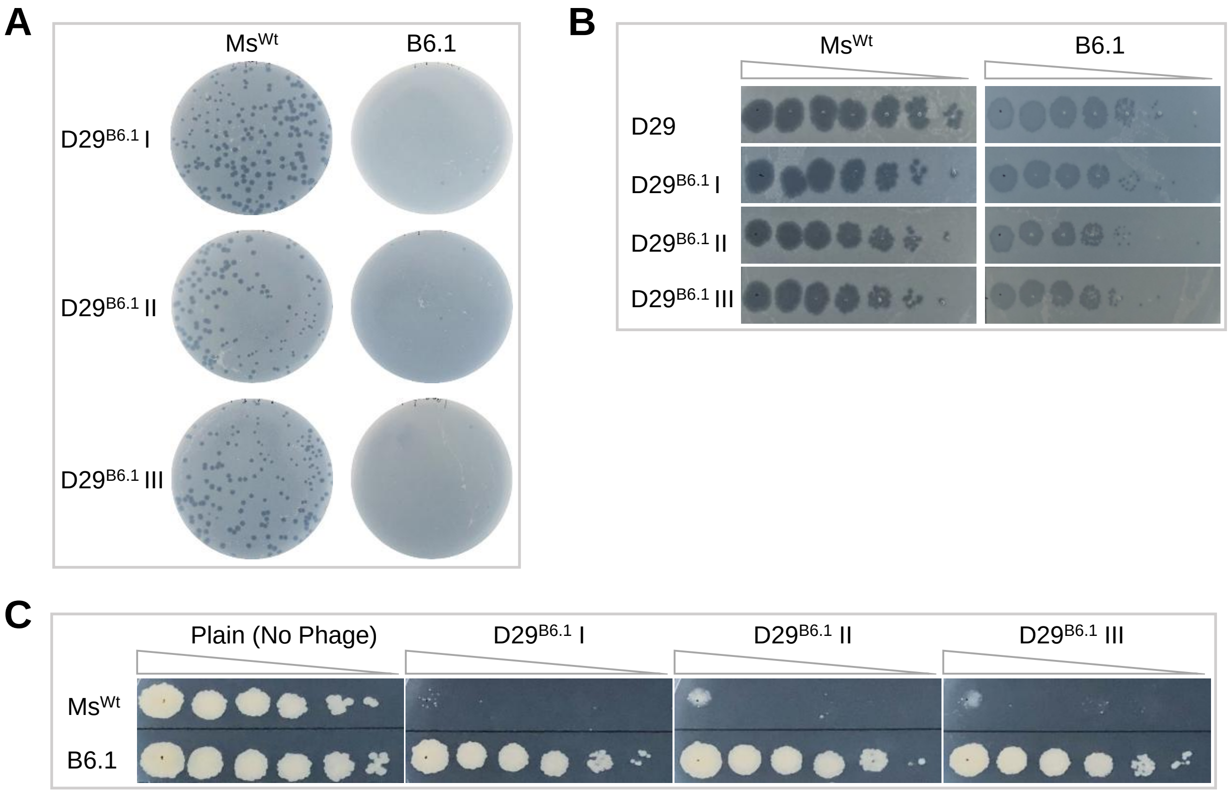


**Figure S5 D29 plaque purification [from] and secondary infection of B6.1.** Primary D29 infection yielded a few plaques on B6.1, three ($D29^{B6.1}$ I, $D29^{B6.1}$ II and $D29^{B6.1}$ III) of which were recovered, amplified and used for secondary infection of B6.1 by plaque assay (A), spot-kill assay (B) and testing on D29-seeded 7H10 plates (C). The susceptibility profile of B6.1 to $D29^{B6.1}$ I, $D29^{B6.1}$ II and $D29^{B6.1}$ III remains consistent with its susceptibility to D29, which is characterized by turbid plaques/zones of growth inhibition.

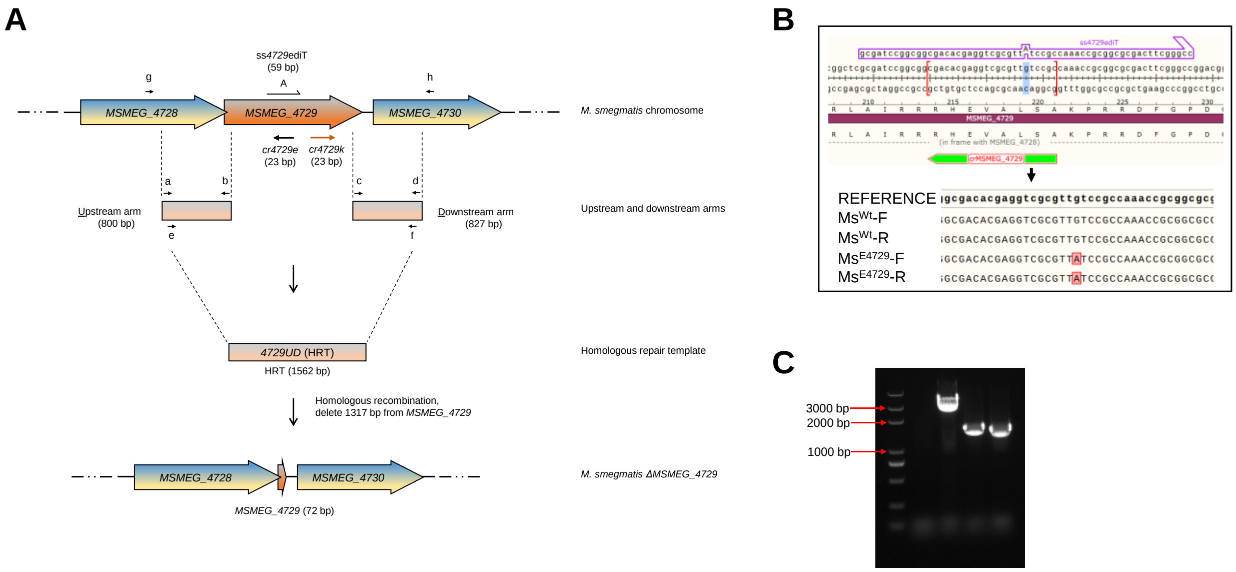


**Figure S6 CRISPR/Cas12a-assisted recombineering of *MSMEG_4729*.** A, Schematic for recombineering in *M. smegmatis*. For the knockout, 2-4 µg each of pCR*4729*k and *4729UD* were transformed into electrocompetent Ms:pJV53-Cpf1 cells by electroporation. Transformants were verified by PCR and Sanger sequencing using the primer pair g/h. For targeted base editing, pCR*4729*e and ss*4729*ediT were transformed into electrocompetent Ms:pJV53-Cpf1 cells as crRNA and HRT respectively. Base editing was verified by Sanger sequencing. B, The *4729G657A* synonymous mutations was introduced by CRISPR/Cpf1-assisted recombineering using the homology-directed repair pathway. Sequences and orientation of the crRNA and the single-stranded editing template as well as the target base are indicated in the upper panel. Base editing was verified by Sanger sequencing of forward (F) and reverse (R) sequences at the site in both wild type ($Ms^{Wt}$) and edited strain ($Ms^{E4729}$). C, PCR verification of *MSMEG_4729* knockout. Lane 1, 2K+ DNA ladder; lane 2, blank control (nuclease-free water); lane 3, negative control ($Ms^{Wt}$, 3019 bp); lanes 4 & 5, knockout strains ($Ms^{\Delta 4729}$, 1702 bp).

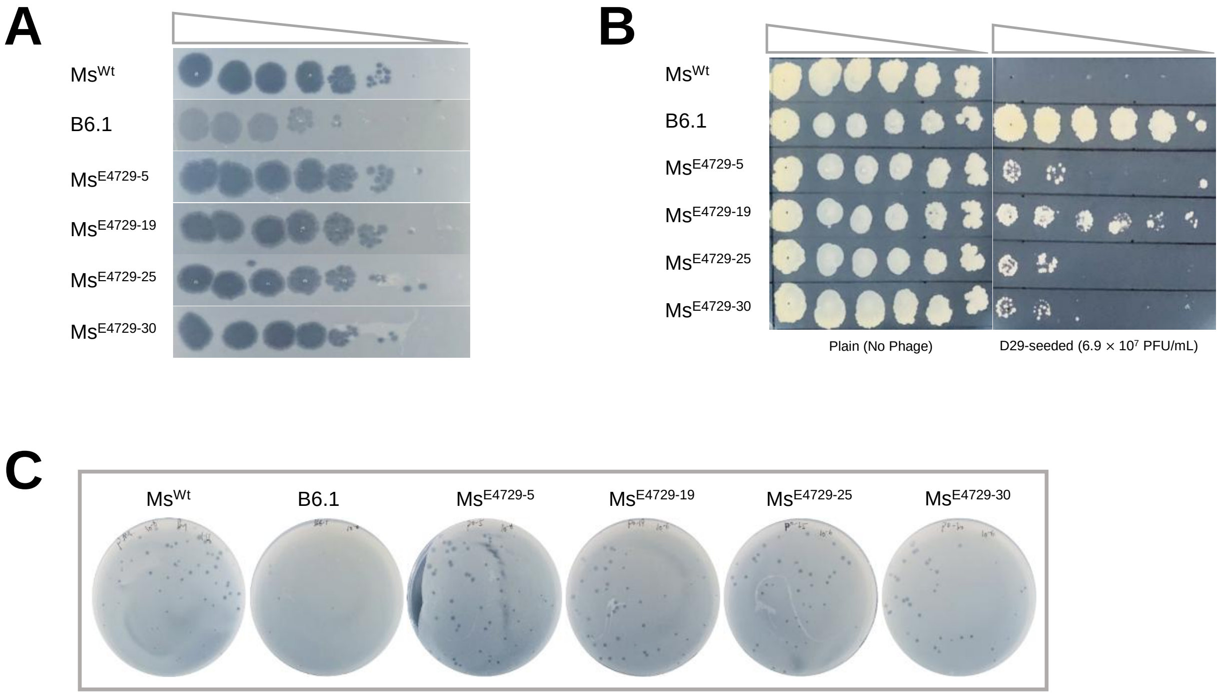


**Figure S7 D29 susceptibility testing of purified edited strains prior to unmarking.** The edited strain $Ms^{E4729}$ was purified by plating of 10-fold serial dilutions, and 30 single colonies were re-verified for the presence *47279G657A* mutation. Four (4) of the colonies (colonies #5, #19, #25 and #30) have the mutation, but spot-kill (A) and plaque (C) assays suggest that their susceptibility to D29 remains similar to that of $Ms^{Wt}$, but differs on D29-seeded 7H10 plates (B).

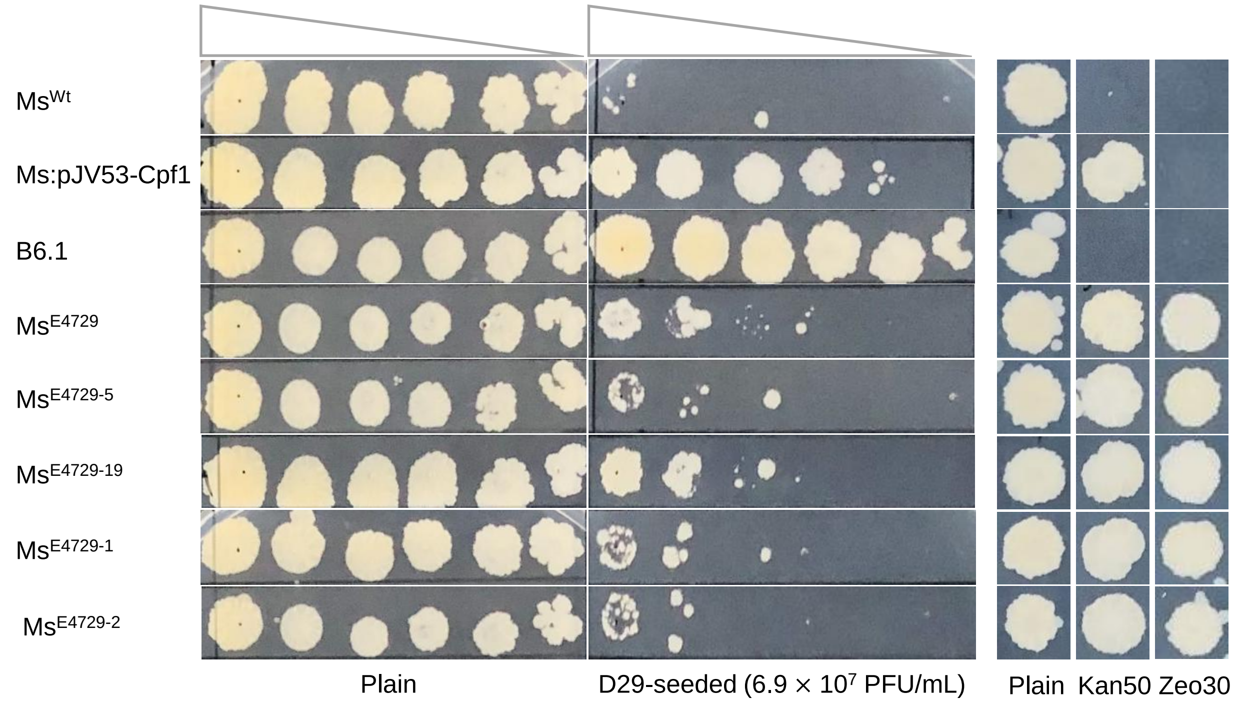


**Figure S8 Verifying possible plasmid role in background bacterial growth on D29-seeded plates.** Base-edited strains manifested similar susceptibility profile as $Ms^{Wt}$ on D29-seeded plates after they were cured of the two recombineering plasmids pJV53-Cpf1 and pCR-Zeo. It appears that *47279G657A* is not responsible for the background resistance of edited strains on D29-seeded plates. This is because Ms:pJV53-Cpf1 as well as $Ms^{E4729-1}$ and $Ms^{E4729-2}$ are also able to grow on D29-seeded plates, and unmarking of edited strains reverses resistance. Ms:pJV53-Cpf1, $Ms^{Wt}$ harboring pJV53-Cpf1; $Ms^{E4729-1}$ and $Ms^{E4729-2}$, colonies #1 and #2 respectively from purification of $Ms^{E4729}$, but do not have the *47279G657A* mutation.

**Table S2 Strain selection for epigenome sequencing**

| Strain | Has mutation? | Has IS transposition? | Selection |
|---|---|---|---|
| $Ms^{Wt}$ | - | - | Selected |
| A12.1 | No | Yes | Not selected |
| A24.3 | Yes [a] | Yes [c] | Not selected |
| A33.1 | No | Yes | Not selected |
| A51.3 | Yes [a] | Yes | Not selected |
| B6.1 | Yes [a] | No | Selected |
| B11.3 | No | No | Selected |
| B15.1 | No | No | Selected |
| B28.2 | Yes [a] | Yes | Not selected |
| B35.1 [b] | No | Yes | Not selected |
| B54.2 | Yes [a] | No | Selected |
| C77.1.1 | No | No | Selected |

[a] This strain has the *4729G657A* synonymous mutation only.

**Table S3 Number of restriction-modification proteins in different *M. smegmatis* strains**

| Strain | Genome size (bp) | Number of R-M proteins |
|---|---|---|
| INHR1 | 6,988,337 | 2 (1M, 1R) |
| INHR2 | 6,988,302 | 2 (1M, 1R) |
| JS623 | 6,464,916 | 10 (6M, 2R, 2S), including 2M on plasmid pMYCSM02 |
| $mc^2$ 155 | 6,988,269 | 2 (1M, 1R) |
| NCTC8159 | 6,983,267 | 6 (2M, 4R) |

M, modification protein; R, restriction protein; S, specificity protein.
M methylates the DNA; R cleaves the DNA; and S specifies the DNA sequences that R and M recognize.

**Table S4 Matched restriction-modification proteins in *M. smegmatis***

| Motif | Gene | Product | Strains |
|---|---|---|---|
| CTCGAG | *MSMEG_3213* | Type II methyltransferase | INHR1, INHR2, $mc^2$ |
| CTYRAG | *MSMEG_3214* | Type II restriction endonuclease | 155, NCTC8159 |

Y = C or T; R = A or G.
